# Chromatin accessibility classification of TAD boundaries discloses new architectural proteins

**DOI:** 10.1101/2024.12.18.629025

**Authors:** Karina Jácome-López, Leonardo Ledesma-Domínguez, Ayerim Esquivel-López, Rosario Pérez-Molina, Andrés Pénagos-Puig, Amaury Aguilar-Lomas, Mayra Furlan-Magaril

## Abstract

3D genome organization is crucial to modulate gene expression. Topologically Associated Domains (TADs) isolate genes and their regulatory elements within the same topological neighborhood, avoiding crosstalk between regulatory elements. Perturbation of domain boundaries causes aberrant genomic contacts and gene expression misregulation. Architectural proteins such as CTCF and the cohesin complex are critical to form boundaries. However, we still lack a complete understanding of what makes a boundary more effective at insulating genomic contacts than others. To understand how domains are structured, we experimentally classified boundaries according to their chromatin accessibility as a proxy of protein occupancy in K562 human cells. We found that highly accessible boundaries are occupied by more proteins, are more robust contact insulators, have a more conserved CTCF DNA-binding motif and are more conserved across cell types in contrast to less accessible ones. By exploring the proteins enriched at boundaries with different accessibility, we found that CTCF and cohesin, together with REST, form a module, and ZFN316, together with EMSY, form another module, and both modules occupy boundaries very frequently. Finally, by using CRISPR-Cas9 mediated genetic edition of ZNF316 DNA-binding motif at a robust domain boundary, we demonstrate that ZNF316 can block chromatin contacts in the absence of CTCF. Our results emphasize the importance of protein combination and abundance to support boundary strength and propose ZNF316 as a novel architectural protein.

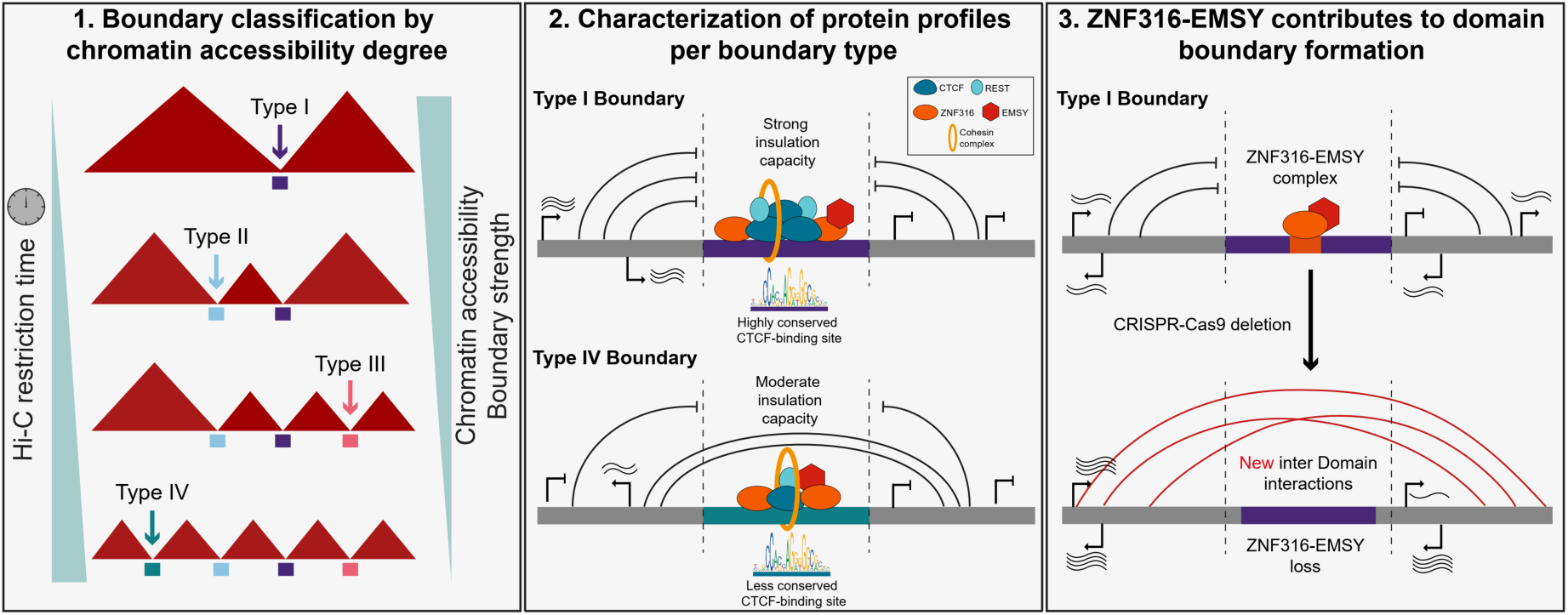

## INTRODUCTION

Topologically Associating Domains (TADs) are structural units that contribute to the spatiotemporal control of gene expression by isolating the interactions between regulatory elements in the genome favoring the interaction between elements inside domains and restricting the frequency of inter-domain interactions (1,2). The isolation of genomic interactions is achieved through the action of domain boundaries that prevent cross-regulation between genomic elements (1–3). Various reports have shown the importance of domain boundaries in maintaining proper gene expression patterns in diverse scenarios including development, cell differentiation and DNA replication, and functional assays have shown boundaries misfunction can lead to pathologies like cancer (4–8).

The human genome contains ∼3,500-8000 domains covering more than 90% of the genome, depending on the resolution used. Their size varies from ∼30-40kb to 3Mb, with a median size of 720-840kb (9,10). 55-75% domains are preserved between human cell lines and 45- 76% are conserved between human and mouse (1,9,11). In addition to mammals, there are other organisms in which domains have been described such as *Drosophila melanogaster* (12), *Schizosaccharomyces pombe* (13), the X chromosome of hermaphroditic *Caenorhabditis elegans* individuals (14), in the bacteria *Caulobacter crescentus* (15) and in *Oryza sativa* (16).

There is a high heterogeneity regarding the amounts of architectural proteins present at boundaries, histone modifications and gene transcription. Domain boundaries are enriched in DNA-binding sites for proteins such as CTCF, cohesin, Mediator, RNA Pol II and general transcription factors (1,17). Also, boundaries are rich in genes that are expressed constitutively and in H3K27ac, H3K4me3, H3K4me1 and H3K36me3 histone modifications and they are also DNase I chromatin accessibility sites (1,9,18). CTCF is a key factor in forming and maintaining domain boundaries and forming chromatin loops together with cohesin (19,20). Chromatin loop extrusion is implicated in loop and domain formation, where the cohesin complex runs through the chromatin fiber until it finds CTCFs bound in a convergent orientation, stabilizing loop anchors or domain boundaries (21–24).

The deletion of specific CTCF DNA-binding sites in boundaries results in gene expression pattern alteration via the incorrect formation of contacts between distal regulatory elements leading to developmental abnormalities (3,4,7,25). Also, CTCF acute depletion causes domain disappearance from most genomic regions with ∼18% of domains remaining intact (19). Nevertheless, there are studies where CTCF depletion does not seem to fully affect boundary function (11,26,27).

Today, there are still many questions about the differences between boundaries in terms of their insulation capacity and their direct influence on transcriptional regulation. Thus, characterizing the identity and protein occupancy levels between domain boundaries is essential to discern whether there is a functional hierarchy in their insulating capacity. Here we have classified domain boundaries by chromatin accessibility as a proxy for protein occupancy. We found that highly accessible chromatin boundaries are occupied by more proteins, are more robust contact insulators, have a more conserved CTCF DNA-binding motif and are more conserved across human cell types. Also, we found a collection of proteins (CTCF, Cohesin, REST, ZNF316 and EMSY, among others) that occupy boundaries. Importantly, CTCF, Cohesin and REST form a module that often binds together at boundaries. In contrast, ZNF316 and EMSY form a module that frequently binds together at boundaries, and the latter module is prominent at boundaries without CTCF. Finally, we functionally demonstrate that ZNF316 DNA-binding motif edition leads to domain boundary disruption and increased inter-domain interactions, suggesting ZNF316 as a new factor structuring chromatin and insulating interactions between regulatory elements.

## MATERIALS AND METHODS

### Cell culture

K562 cell line were obtained from ATCC (ATCC® CCL-243™), the cells were grown in Iscove’s Modified Dulbecco’s Medium (IMDM) supplemented with 10% BFS and maintained in 5% CO_2_ at 37°C.

### Hi-C in nuclei experiments

Hi-C at different times of digestion enzyme were performed with 2 million of cells in two replicates. The fixation occurred during 10 minutes at 2% formaldehyde (Pierce™ 16% Formaldehyde, Methanol-free Thermo Scientific™) in supplemental medium. After that, glycine was added to 0.13M final concentration and incubated first for 5 minutes at room temperature and then for 15 minutes on ice. The pellets were washed with cold PBS buffer. The cell lysis was performed in lysis buffer:10mM Tris-HCl pH8, Igepal 0.2%, NaCI 10mM and complete protease inhibitor at 1X (Sigma-Aldrich®) during 30 minutes on iced. Then the pellets were washed twice, with lysis buffer and then with New England Buffer #2 (NEB2) at 1.25X concentration.

The pellets were incubated in 360μl of NEB2 1.25X, 11μl of SDS 10% incubated at 37°C for 45 minutes with agitation 950rpm. Then 75μl 10% Triton were added under the same incubated conditions as before. The digestion occurred with 1500U *HindIII* (New England Biolabs®) and incubated in different time points: 0.25, 1, 3 and 12 hours at 37°C 950rpm. The fill-in was performed by adding 1.5μl of the nucleosides dCTP, dGTP and dTTP each of them at 10mM, 37.5μl Biotin-14-dATP (Invitrogen™) 0.4mM and (10μl) 50U Klenow enzyme DNA pol I large fragment (New England Biolabs®) . The fill-in reactions were incubated at 37°C for 75 minutes in agitation (700rpm) during 15s each 30s.

To each experiment 100μl 10X ligation buffer (New England Biolabs®), 10μl BSA (10mg/ml), 25U T4 DNA ligase (Invitrogen™) and 352μl H_2_O were added. The ligation reactions were incubated overnight at 16°C.

For reverse crosslinking we added 50μl proteinase K (10mg/ml) and 10μl RNase (10mg/ml) and purified with *Phenol*/*Chloroform*/*Isoamyl Alcohol* 25:24:1 (Invitrogen™). The DNA was sonicated in COVARIS S220 (Covaris, Inc.) with the following conditions: Duty factor 10%, Peak incident power (w) 140, Cycles per Burst 200 and H_2_O level 10-15, during 80 seconds. The biotin remove and extreme repair were performed using 7µg of sonicated DNA in a volume of 130µl TLE, 16μl 10X ligation buffer NEB, 2μl dATP 10mM, 15U T4 DNA polymerase NEB (New England Biolabs®) y 7μl H_2_O were added and incubated 30min at 20°C. Then we added 5μl dNTPs 10mM, 4μl 10X ligation buffer NEB, 5μl T4 Polynucleotide Kinase (New England Biolabs®) (10U/μl), 1μl Klenow large fragment (New England Biolabs®) (5U/μl) and 25μl de H_2_O, this reaction was incubated for 30 minutes at 20°C. The DNA were selected using Ampure SPRI Beads (Beckman Coulter, Inc.) to get 300-600bp fragments. The pull-down was performed with 150μl Dynabeads MyOne Streptavidin C1 (Invitrogen™). The libraries were marked with TrueSeq DNA Adapters (Illumina, Inc.) and amplified using Phusion High-Fidelity DNA Polymerase (New England Biolabs®). The paired-end sequencing was performed in a HiSeq2500 (Illumina, Inc).

The Hi-C experiments in the cells edited by CRISPR-Cas9 were performed as described above, but with the following modifications: Per experiment 5 millions of cells were used. The chromatin digestion we used 400U of DpnII enzyme (New England Biolabs®) incubating overnight. The paired-end sequencing was performed in Illumina NovaSeq 6000 (Illumina, Inc).

### Restriction percentage of chromatin digestion

The chromatin temporal digestion and DNA extraction were performed as described in Hi-C experiments section. The evaluation of digestion percentage at chosen boundaries was performed with primers that flank a *HindIII* site. The qPCR reactions were made with 50ng of DNA using the KAPA SYBR Fast qPCR kit (Kapa Biosystems). The restriction percentage was obtained with the formula: % 𝑅𝑒𝑠𝑡𝑟𝑖𝑐𝑡𝑖𝑜𝑛 = 100 – 100/2˄[(𝐶𝑡𝑅–𝐶𝑡𝐶) Digested – (𝐶𝑡𝑅– 𝐶𝑡𝐶) Undigested] (28).

### CRISPR-Cas9 mediated edition

Two sgRNAs were designing in CRISPOR (29). The sgRNAs were cloned in the lentiCRISPRv2 (Addgene) vector based on [lentiGuide oligo cloning addgene protocol](https://media.addgene.org/data/plasmids/52/52961/52961-attachment_B3xTwla0bkYD.pdf). Lipofectamine 3000 (Invitrogen™) was used to transfect 3X10^5^ cells with 2µg of vector (sgRNA-1 and sgRNA-2). The transfected cells were selected with puromicine. Clonal dilutions were made to isolate cells, we evaluated the deletions by end-point PCRs and sequencing. See sgRNAs sequences in Supplemental Table S1.

### RT-qPCR

Total RNAs were isolated using Invitrogen TRIzol^TM^ reagent as user guide recommend. The RNAs were treated with RQ1 RNase-Free DNase (Promega). The cDNAs synthesis were performed with SCRIPT cDNA Synthesis Kit (Jena Bioscience). The OAZ1 gene was used as housekeeping gene calibrator. The reactions were performed in the Applied Biosystem StepOne Real-Time PCR System using KAPA SYBR Fast qPCR kit (Kapa Biosystems). All primers used are available in Supplemental Table S1.

### Hi-C data analysis

The paired-end raw fastq files were processed in HiC-Pro v2.11.1 with default parameters (30). The alignment of the reads was performed with Bowtie2 v2.3.4.1 against the hg19 human genome. The HiC-Pro quality output files were visualized in MultiQC v1.7. The normalized ICE matrices were generated with same number of valid pairs and computed at 50kb resolution for the Hi-Cs at different digestion time points at 20kb and for the Hi-Cs in the edited and control cells. The Hi-C replicates at each time point showed a similar QC profile (Supplemental Table S2). Also, the number of Hi-C contacts generated between regions at different genomic distances was plotted to verify that the *cis* decay was equivalent between replicates (Supplemental Figure S1). Finally, the matrices were visually inspected to confirm similarities between replicates. Afterwards all analysis was performed in the merged data sets of the replicates. The quality metrics for all Hi-C experiments presented are in Supplemental Table S2.

### TAD calling and boundary accessibility classification

The domains were called with the measure of TAD-separation score in HiCExplorer (31) at 50kb resolution. The comparisons were made with False Discovery Rate (FDR) and q-value threshold of 0.001. The domains and boundaries were aggregated by their identification at each Hi-C time point using the exact coordinates. The insulation scores of boundaries were computed in TADtool with a window size of 100,000bp and a TAD cutoff of 15 (32). The corner scores of the domains were extracted from published data in K562. (9). The TAD boundaries identified in this study are in Supplemental Table S3. Inter Domain Contacts or Inter-TAD score represents the number of interactions that cross a boundary. These values were obtained at 50kb of resolution with a window of 150kb. The inter-TAD value was calculated as described (33).

### Boundaries and domains conservation across human cell types

Published Hi-C data sets for the cell lines GM12878, HUVEC, HMEC, IMR90, NHEK, K562 and HCT116 were used (See Data Availability). HiCExplorer was used to identify domain boundaries as described above. The conserved boundaries across the different cell types were identified using a window +/- 125kb from the start and end coordinate of the boundaries. Conserved domains between cell line types were identified using an interval of +/- 125kb from their start and end coordinates. To be considered a conserved domain, the overlap must be at least 80%. The conserved boundaries and domains were identified using searchV2.1.pl (Supplemental material).

### Fast-ATAC seq processing data

Published Fast-ATAC seq data were downloaded (Data Availability) (34). The fastq files were aligned with HISAT v2.1.0 against the hg19 genome version. The peaks were called with Genrich v0.6 removing the PCR duplicates and applying the default parameters. Peaks larger than 1kb were removed. The read coverage was computed with multiBamSummary in deepTools v3.5.0 (35).

### A/B compartments analysis

The PCA eigenvalues were computed with hicPCA in HiCExplorer for Hi-C matrices at 50kb of resolution using the Lieberman parameter. H3K27ac and H3K4me3 histone modifications from ENCODE mapped to the hg19 genome were used to assign the active regions (see Data Availability).

### ChIP-seq data analysis

296 ChIP-seq data sets produced by ENCODE in K562 cell line were used (see Data availability). We analyzed the optimal IDR thresholded peaks files processed based on the ENCODE processing pipeline using the hg19 genome. The identification of peaks located at boundaries were made in bedtools v2.29.0. The average signal score for CTCF, REST, ZNF316 and EMSY were made using bigwig files and processed with multiBigwigSummary in deepTools v3.5.0. The boundary classifications, with and without CTCF, was performed based in three different CTCF ChIP-seq sets (see Data availability).

### RNA-seq data analysis

RNA-seq data sets produced by ENCODE in K562 cell line were used (see Data availability). The alignments bam files of two isogenic replicates processed using the hg19 genome version were analyzed with multiBamSummary in deepTools v3.5.0. The gene quantification files from same data set were analyzed to identify the gene expression level at boundaries.

### Gene Ontology enrichment

The gene ontology enrichment analyses were made in gProfiler (36). The default parameters were applied. The statistical domain scope used was the option all known genes.

### DNA-binding motifs orientation analysis

The CTCF DNA-binding motifs were identified using the FIMO tool from MEME suite (37). The CTCF sequence motif used was the reported in HOMOCO V11 database. The CTCF motifs were identified based on the DNA sequence from the ChIP-seq peaks. The extraction of motifs inside the boundaries was made with bedtools intersect. The boundaries of a total size of 50kb were divided in 10kb windows to map the strand motif orientation in forward and reverse. For each window, the frequency was calculated dividing the number of motifs, forward or reverse, between the total number of motifs for each type of boundaries. The same process was made for REST DNA-binding motif. For ZNF316, we processed the fasta ChIP-seq peak sequences to find a consensus motif with MEME suite, we used the HOMOCO V11 database to find the best possible motif sequence match. The DNA-binding motif orientation analysis was performed as described above for CTCF.

### CTCF DNA-binding motif conservation analysis

The CTCF DNA sequence motif conservation was extracted from phasCons46way database. The motifs sequences were extended to 60bp. The conservation score graphed corresponds to the mean of all nucleotides scores for each position.

## RESULTS

### The degree of chromatin accessibility at domain boundaries reflects their insulation capacity

The structure and maintenance of chromatin domains are presumably due to the proteins occupying their boundaries (1,18,38,39). We hypothesized that more robust boundaries have higher chromatin accessibility and therefore higher protein occupancy. Thus, domains with more chromatin-accessible boundaries should be more stable than those with fewer.

To determine if accessibility indeed reflects more robust boundaries and domains, we experimentally explored whether domain boundaries exhibit different degrees of accessibility in the human K562 cell line. To do so, we performed Hi-C experiments varying the digestion time with a restriction enzyme. Restriction enzymes have been used to measure chromatin accessibility in different contexts before, as more accessible chromatin will be digested faster (40–43). The incorporation of this digestion time course into the Hi-C enables us to directly assess boundary accessibility and TAD formation from the same experiments (Fig. 1a). We digested chromatin for 0.25, 1, 3 and 12 hours to capture the most accessible boundaries at the shortest incubation times. The results show that chromatin is indeed digested gradually over time. Chromatin digestion is readily observed from 0.25 hours of incubation and the percentage of digestion increases with longer digestion times (Fig. 1b). After digestion, we continued the Hi-C experiment and sequenced the generated libraries with good quality in terms of valid pairs and *cis:trans* contact ratios (Fig. S1a-b). We next built contact matrices from the different time points and called TAD boundaries using TAD-separation score. The number of domains detected increased with the digestion time (Fig. 1c). Next, we classified boundaries based on their accessibility. The boundaries identified from 0.25 hours of digestion are those with the highest accessibility, and we grouped them as type I. Boundaries identified from one hour of digestion are type II, boundaries identified from three hours are type III. Finally, boundaries identified only after twelve hours of digestion are type IV, representing the least accessible boundaries detected (Fig. 1d and S1c).

**Figure 1.**
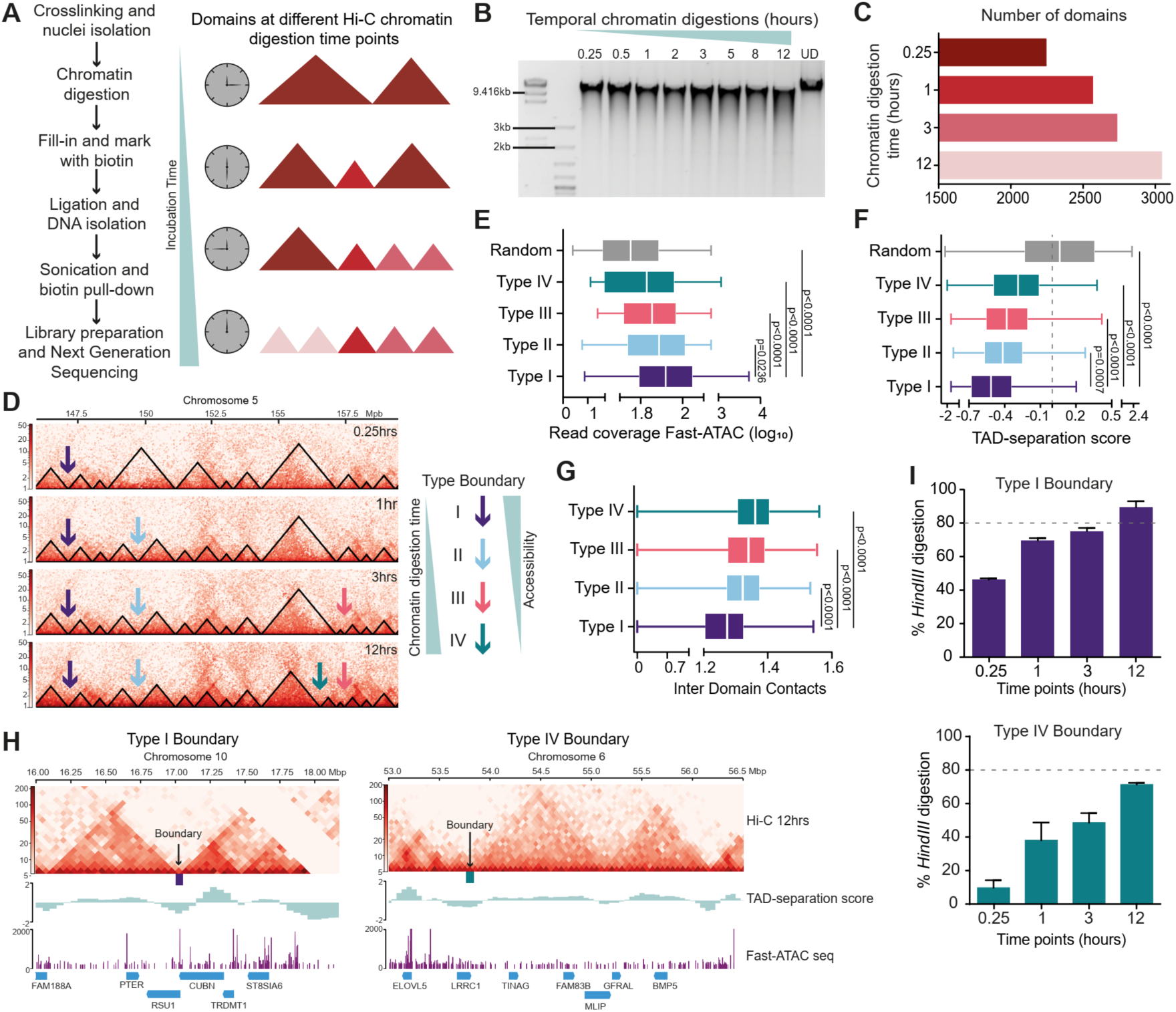
Chromatin accessibility reflects insulation capacity of domain boundaries. (A) Hi-C experimental procedure incubating the *HindIII* restriction enzyme at different time points, recovering the total number of domain boundaries in the longest incubation time. (B) Temporal digestion with *HindIII* since 0.25 to 12 hours, demonstrating the gradual chromatin digestion. As a control an undigested chromatin sample is shown (UD). (C) The Hi-C assays were performed at four time points: 0.25, 1, 3 and 12 hours. The number of domains and boundaries increased proportionally to the time. (D) Examples of the Hi-C matrices derived from the different restriction timepoints. The domain boundaries identified were classified as type I, reflecting the most accessible boundaries. The type II, II and IV boundaries were recognized from 1 hour onwards, 3 hours onwards and 12 hours, respectively. The type IV boundaries are the least accessible only appearing at 12 hours. (E) Fast-ATAC seq read counts in the different boundaries’ type. The type I boundaries, classified as most accessible, showed the higher count of Fast-ATACseq reads. As a control, the read count was made in random genomic regions of 50kb size. Kruskal-Wallis test was applied followed by Dunn’s post hoc test. (F) The TAD-separation score was employed to call domain boundaries; the lowest scores represent the boundaries with enhanced insulation capacity. Kruskal-Wallis test was applied followed by Dunn’s post hoc test. (G) The inter-TAD values were computed, representing the number of interactions that are crossing a boundary. The type I boundaries allow a low number of interactions between domains, in contrast the type IV boundaries allow more inter domain contacts. Kruskal-Wallis test was applied followed by Dunn’s post hoc test. (H) Example of type I and type IV boundary. The Hi-C matrices represent 12 hours of *HindIII* incubation, below the TAD-separation score is plotted and the Fast-ATACseq displayed. (I) Chromatin digestion in the boundaries represented in (H). We designed primers flanking restriction sites for *HindIII* in the boundary and calculate the percent of digestion through qPCR. The percent of digestion in this type IV boundary in lower comparing with the type I boundary.

Using available Fast-ATAC seq data, we assessed the accessibility degree of all boundaries. The results confirmed that our experiments with different restriction incubation times recapitulate different levels of boundary accessibility measured through ATAC, with type I boundaries having the most Fast-ATAC seq signal and type IV boundaries having the least Fast-ATAC seq signal (Fig. 1e).

To analyze how accessibility relates to boundary insulation capacity, we computed the TAD- separation score for all boundaries. The TAD-separation score of boundaries with higher accessibility displays more negative values than boundaries with less accessibility, reflecting they are more robust at insulating interactions from adjacent regions (Fig. 1f). This observation was corroborated by counting the inter-domain contacts (Inter-TAD) (Fig. S1d). We found that the lower the accessibility, the more contacts cross the boundary (Fig. 1g,e). Finally, to further confirm our finding, we detected boundaries using the Insulation Score (IS) (32, 44). We found that type I boundaries have lower IS values, reflecting fewer interactions across them compared to type IV boundaries (Fig. S1e).

Next, we mapped our domain boundaries and domains to a public and deeply sequenced Hi-C data set at 5kb resolution (9). We obtained their corner score and calculated their TAD- separation score. Using 5kb boundaries, we confirmed that the more accessible type I boundaries are more robust, with higher corner scores and lower separation scores than less accessible type IV boundaries (Fig. S1f). This confirms that our boundary detection, accessibility classification and conclusions are sustained using a higher-resolution Hi-C data set, reducing the boundaries to 5kb.

To validate our genome-wide observations experimentally and locally, we explored the accessibility of selected type I and IV boundaries. To do so, we measured the digestion percentage of amplicons containing the boundaries, using the chromatin digested at different time points as a template for PCR reactions. The results show that the selected type I boundaries have higher restriction percentages at earlier time points than type IV boundaries, confirming they are more accessible (Fig. 1h-i). Additional type I and IV boundaries were tested, confirming that type I boundaries are more accessible (and digested first) than type IV boundaries (S2a-d).

### The orientation and conservation of CTCF DNA-binding motifs influence boundary strength

We next mapped the different types of boundaries to A and B compartments. We defined the compartments through a Principal Component Analysis (PCA) using as a seed the signal for the H3K27ac histone mark to assign the A compartment. Each genomic window has a positive or negative PC1 value and in our analysis the positive values are directly related to the H3K27ac enrichment.

Unexpectedly, only 54% of type I boundaries are in the A compartment, a percentage that is similar for all boundary types (Fig. 2a). When plotting the PC1 values, we observed that type I-III boundaries have greater enrichment of H3K27ac than type IV boundaries (Fig. 2b). This is also observed when using H3K4me3 histone mark as the seed (Fig. S3a). These observations suggest that compartment type does not reflect the degree of chromatin accessibility by protein occupancy, since together the time course of digestion in the Hi-C, the public ATAC data and our validation of the accessibility of the percentage restriction analysis, show that the boundaries have different levels of accessibility. However, the acetylation levels at type I boundaries are significantly higher than at type IV boundaries.

**Figure 2.**
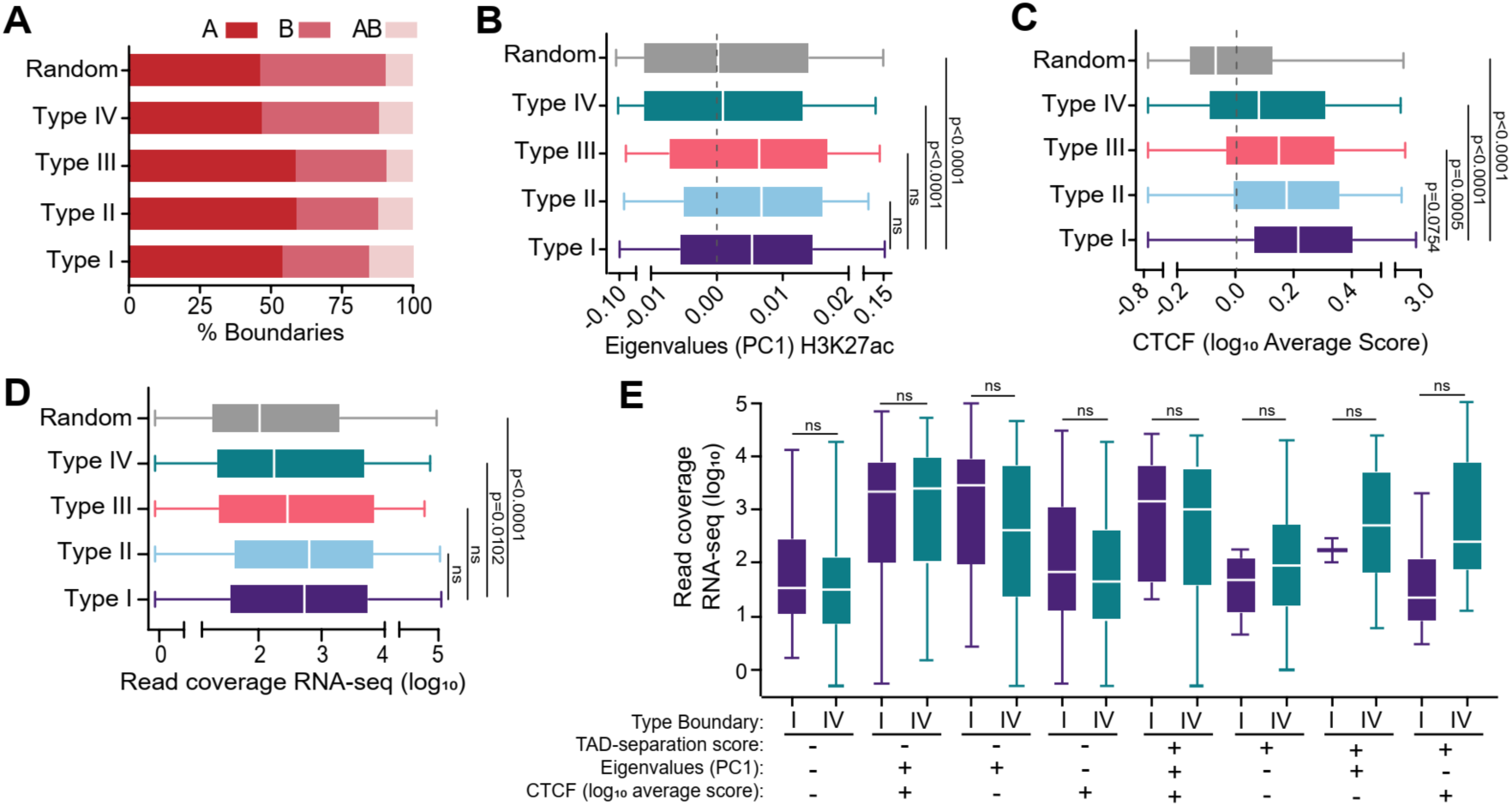
The accessibility degree of boundaries correlates with more CTCF and positive PC1 values. (A) Percent of boundaries located in compartment A, B or AB. (B) PC1 values to call compartments, using H3K27ac histone mark as a seed. Kruskal-Wallis test was applied followed by Dunn’s post hoc test. (C) CTCF average signal in the four types of boundaries along the 50kb. Kruskal-Wallis test was applied followed by Dunn’s post hoc test. (D) RNA-seq read coverage at the different boundary types. Random regions of 50kb were measure. Kruskal-Wallis test was applied followed by Dunn’s post hoc test. (E) RNA-seq read coverage for boundaries type I and type IV with the same values in TAD-separation score, PC1 values and CTCF. Mann-Whitney statistic test report no significant difference between boundary types.

It is well documented that CTCF is enriched at domain boundaries, which contributes to their formation together with the cohesin complex, as detailed above (20–24). We assessed CTCF abundance at our different boundary types using available ChIP-seq data (see materials and methods). The average signal enrichment of CTCF protein within the boundary was calculated, and we discovered that type I boundaries have a higher average signal enrichment than type IV boundaries (Fig. 2c). These results show that CTCF concentration at boundaries is partly responsible for increasing their accessibility.

To better understand the complexity of type I and IV boundaries, we analyzed together the presence of CTCF, the TAD-separation score and the PC1 compartment values generated using the H3K27ac histone mark as the seed. By looking only at two variables at a time, we observed that 60% of type I boundaries have a negative TAD-separation score and are in compartment A, while only 48% of type IV boundaries have these characteristics (Fig. S3b). The 81% of type I boundaries have CTCF enrichment with negative TAD-separation scores values, in contrast to 51% of type IV boundaries (Fig. S3c). When analyzing the presence of CTCF and the A compartment assignment we found that 55% and 43% of type I and IV boundaries show a correlation between the presence of CTCF and A compartment, respectively (Fig. S3d).

When analyzing these three characteristics together, we note that 54% of type I and 37% of type IV boundaries show negative TAD-separation score values, are in compartment A and are enriched with CTCF (Fig. S3e). The second most representative category of type I boundaries, which is 27%, exhibits negative TAD-separation score values, B compartment and CTCF enrichment. Notably, 26% of type IV boundaries show negative TAD-separation score values, are in compartment B, and do not have CTCF enrichment. There is a higher correlation of CTCF with compartment A in both type I and IV boundaries.

We then assessed the coverage of RNAs at boundaries. Type IV boundaries show less transcripts than type I boundaries (Fig. 2d). We then compared type I and IV boundaries with the same characteristics regarding TAD-separation score, compartment assignment and CTCF signal. The results show no significant differences in transcript accumulation between type I and IV boundaries with the same attributes (Fig. 2e). These results suggest that the combination of CTCF concentration, compartment assignment and transcript levels, make a boundary more robust in structuring domains even though there are fewer type IV boundaries with all three characteristics.

As mentioned above, CTCF is enriched at domain boundaries and several reports have shown that domains disappear upon acute CTCF depletion (19). In addition, the removal of CTCF DNA-binding sites results in boundary disruption and deregulation of gene transcription (6,7,45,46). However, there are some regions where the absence of CTCF does not alter the domain structure (11,26,27). Also, there are boundaries that lack CTCF (47). Our results show that there is a higher proportion of boundaries that have CTCF and are located in the A compartment. However, these features are observed for type I and IV boundaries, so the specific function of CTCF in boundary strength is not clear as type I boundaries are more robust than type IV boundaries.

To better understand the function of CTCF in boundary strength, we investigated in more detail CTCF occupancy at type I and IV boundaries. Using available ChIP-seq data, we identified the boundaries that have one or more CTCF peaks and those boundaries that lack CTCF. Most boundaries exhibit at least one peak of CTCF, but 11% and 26% of type I and IV boundaries do not have CTCF, respectively (Fig. 3a). There is no significant difference in accessibility measured by Fast-ATAC seq between type I boundaries with or without CTCF, which is consistent with our boundary classification based on chromatin accessibility. However, the subset of type IV boundaries without CTCF, represents the group with the lowest accessibility (Fig. 3b). The presence or absence of CTCF in the boundaries is also not related to more negative TAD-separation scores; therefore, CTCF does not determine boundary strength when comparing boundaries with the same accessibility degree (Fig. 3c). However, all type I boundaries are more efficient in blocking interactions across domains, regardless of the presence of CTCF (Fig. 3d). Taken together, our results suggest that in addition to CTCF other unknown factors can contribute to form domain boundary. Also, for the boundaries occupied by CTCF, additional factors could make some boundaries more robust than others.

**Figure 3.**
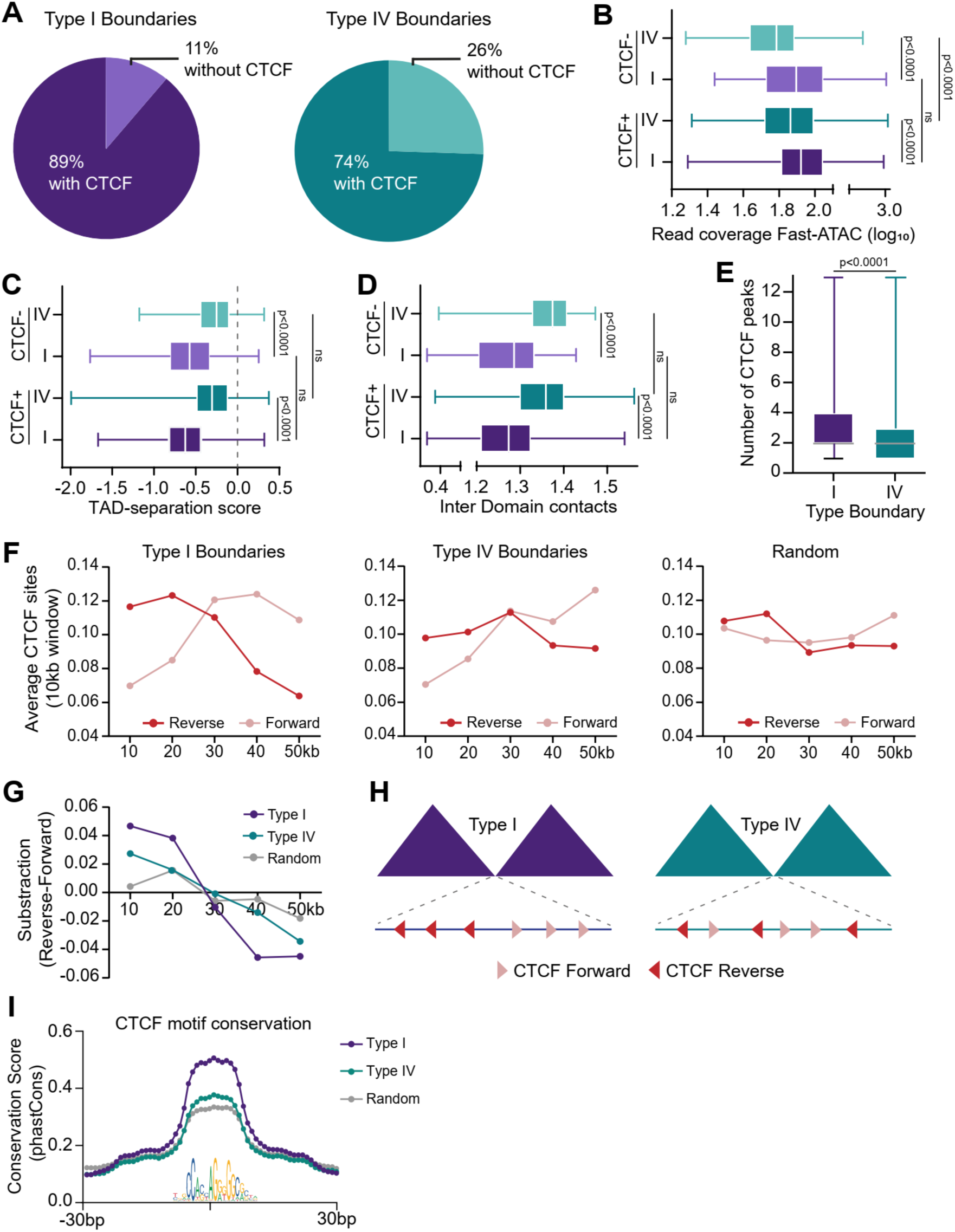
The divergent orientation and the conservation of CTCF DNA-binding motifs drive boundaries’ strength. (A) Percentage of type I and type IV occupied CTCF. (B) Fast-ATAC seq read coverage for boundaries occupied and not occupied by CTCF. Kruskal-Wallis test was applied followed by Dunn’s post hoc test. The accessibility between type I boundaries with and without CTCF showed no statistical difference. In contrast, the type IV boundaries with and without CTCF have statistical difference. (C) TAD-separation scores for type I and type IV boundaries with and without CTCF. Kruskal-Wallis test was applied followed by Dunn’s post hoc test, reporting no significant difference between the same type of boundaries. (D) Inter Domain contact values for type I and type IV boundaries with and without CTCF. Kruskal-Wallis test was applied followed by Dunn’s post hoc test, reporting no significant difference between the same type of boundaries. (E) Number of CTCF peaks along the boundaries. Mann-Whitney test was applied (F) Average of CTCF DNA-binding motif in the forward and reverse orientation along the 50kb boundaries in each 10kb window. The left graph corresponds to type I boundaries, the center graph corresponds to type IV boundaries and the right panel corresponds to random 50kb regions. The red points represent the reverse orientation motif, while the green points correspond to the forward orientation motif. (G) Subtraction of the average values of CTCF DNA-binding motifs in reverse and forward orientation each 10kb window along the 50kb boundaries. The left part of type I boundaries had a predominant motif orientation in reverse, while the right part had a predominant motif orientation in forward. In the type IV boundaries, the divergent profile decreases, while in randomly 50kb regions, there was no evidence of a particular CTCF DNA-binding motif orientation. (H) Diagram of the arrangement of the DNA CTCF binding motif orientation for type I and type IV boundaries (I) Conservation of the CTCF DNA-binding motif for type I and type IV boundaries. Conservation within the type I boundaries is greater than that within the type IV boundaries.

Next, to investigate CTCF abundance at boundaries in more detail, we counted the number of CTCF peaks in the domain boundaries. When comparing the number of CTCF peaks between type I and type IV boundaries, we found significantly more peaks at the type I boundaries. However, both types of boundaries exhibit from one to thirteen CTCF peaks (Fig. 3e).

To determine whether the number of CTCF peaks drives boundary strength, several features including TAD-separation score, PC1 values, RNA read coverage and the number of gene promoters comparing type I and IV boundaries with 0, 1, 2, 3 and so on number of CTCF peaks was analyzed (Fig. S4a). The number of CTCF peaks does not influence the TAD-separation scores. Nevertheless, a clear difference between type I and IV boundaries confirms that the more accessible boundaries are more robust insulators than the least accessible ones (Fig. S4b). Also, the Fast-ATAC reads are similar regardless of the number of CTCF peaks; however, type I boundaries display more reads than type IV (Fig. S4c). The number of CTCF peaks at a boundary, however, does correlate with more positive PC1 values (compartment A) (Fig. S4d), higher RNA read coverage (Fig. S4e), and a greater number of gene promoters (Fig. S4f). This profile is similar between type I and type IV boundaries.

Next, we explored the direction of CTCF DNA-binding motifs at type I and type IV boundaries to determine whether the directionality of CTCF binding is linked to boundary strength. It has been documented that the orientation of CTCF motifs is relevant to form chromatin loops and domains, as the convergent orientation of CTCF binding blocks cohesin chromatin extrusion (21,22). Other studies have shown that inversion or disturbance of CTCF DNA-binding motifs orientation results in decreased insulating capacity of the altered boundaries (47,48,49). We identified the direction of CTCF motifs at type I and IV boundaries. Counting in 10kb genomic windows along the 50kb boundaries and then plotting the frequency and orientation of the motifs. The initial regions of type I boundaries (the first 25kb of the total size of 50kb) show a higher frequency of motifs in the reverse orientation while the final sections of the boundary show a higher frequency of motifs in the forward orientation. Type IV boundaries seem to have an even distribution of the CTCF motif orientation (Fig. 3f-h, S5a-c).

The CTCF motif is conserved between species (50). We next analyzed the conservation level of the motifs embedded in the type I and type IV boundaries based on the vertebrate conservation scores. We found that the CTCF motifs of type I boundaries are more conserved than the CTCF motifs of type IV boundaries (Fig. 3i). This result suggests that it might be CTCF binding affinity or residence time in a more conserved DNA-binding motif that contributes to the boundary strength together with the motif orientation.

### TAD boundaries are occupied by CTCF, Cohesin, REST, ZNF316 and EMSY, among other proteins

Several features associated with domain boundaries have been described. As mentioned, one of the main characteristics is the presence of CTCF and cohesin. In addition, there are reports that proteins such as ZNF143, YY1, and RNA pol II and open chromatin histone marks are also associated with domain boundaries (1,18,51).

To further explore the proteome enriched at domain boundaries we used public ChIP-seq data for 296 proteins in K562 cells and identified proteins occupying boundaries genome- wide (Data Availability, Supplemental Table S4). We first count the total number of proteins occupying boundaries. Type I boundaries have more protein peaks than type IV, which confirms that accessibility is a good proxy for protein occupancy (Fig. 4a).

**Figure 4.**
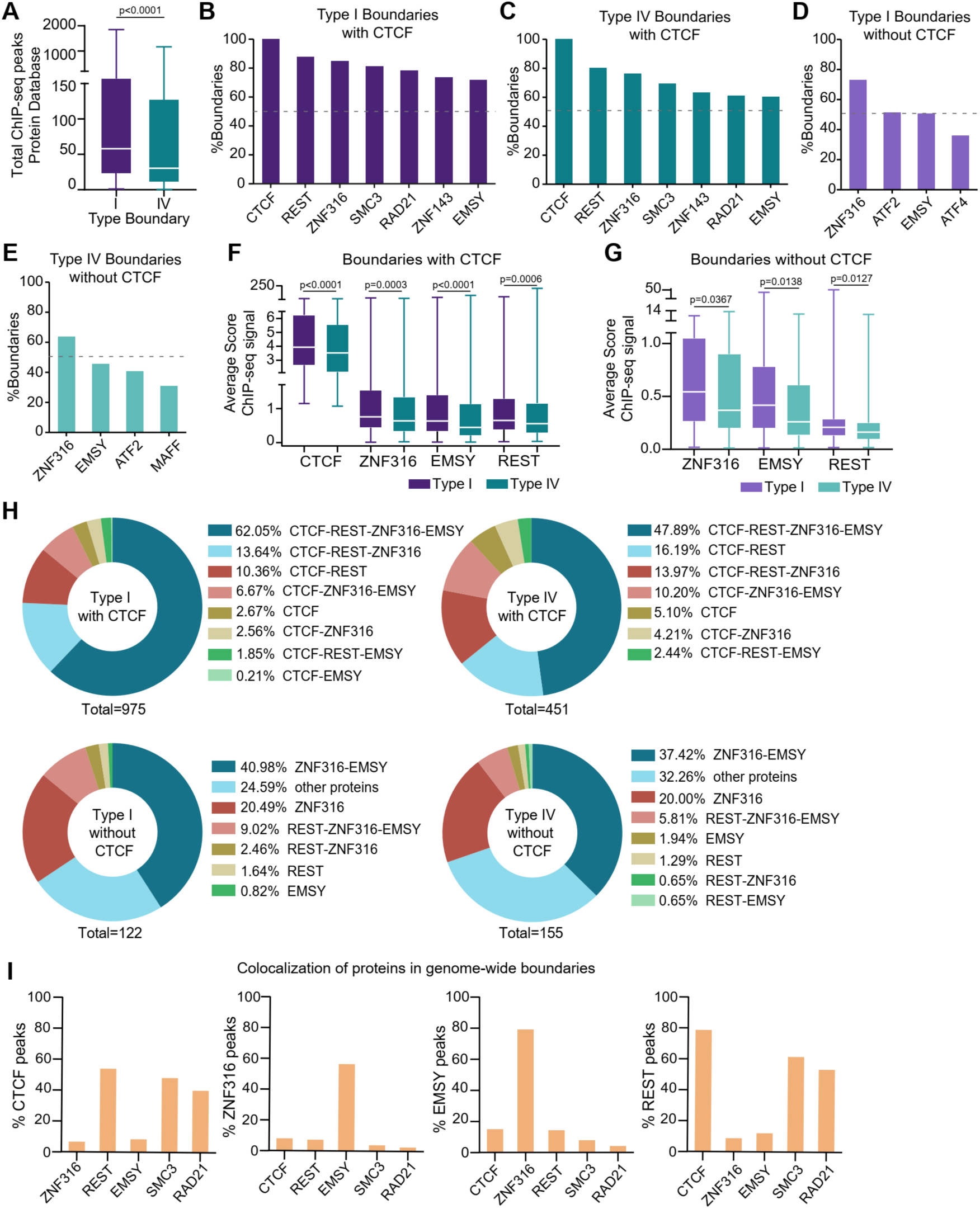
Different proteins including ZNF316, EMSY, REST, CTCF and the cohesin complex occupy domain boundaries. (A) Total protein peaks occupying type I and IV boundaries based on 296 ChIP-seq data sets (see methods). Mann-Whitney test was applied. (B) Percentage of type I boundaries occupied by the 6 most abundant proteins. (C) Percentage of type IV boundaries occupied by the 6 most abundant proteins. (D) Percentage of type I boundaries without CTCF occupied by the 4 most abundant proteins. ZNF316 is present in more than 60% of this subset of boundaries. (E) Percentage of type IV boundaries without CTCF occupied by the 4 most abundant proteins. As in (D), the protein ZNF316 is present in more than 60% of this subset of boundaries. (F) Enrichment of CTCF, ZNF316, REST, and EMSY at boundaries. (G) Enrichment of ZNF316, REST and EMSY at boundaries without CTCF. (H) Classification of domain boundaries based on the presence of CTCF, ZNF316, REST, and EMSY proteins along the 50kb region. The panels above correspond to the boundaries with CTCF, while the panels below correspond to the boundaries without CTCF. (I) ChIP-seq peaks co-localization for CTCF, ZNF316, REST, and EMSY proteins within domain boundaries.

To identify which proteins are enriched at boundaries we considered the 3060 boundaries detected in the overnight digestion condition. As expected, CTCF is found in most boundaries (81%) (Fig. S6a). However, in these boundaries, other proteins are also present (99.5%) (Fig. S6b).

After CTCF, we found that the proteins that are most represented at boundaries genome-wide are REST (*RE1 Silencing Transcription Factor*), followed by the zing finger protein ZNF316, the cohesin complex subunits SMC3 and RAD21, ZNF143 and EMSY (*BRCA2 Interacting Transcriptional Repressor*) (Fig. S6c). Interestingly, boundaries without CTCF are occupied mainly by ZNF316, suggesting a potential function of this protein to preserve a boundary in the absence of CTCF (Fig. S6d). We then measured the proportion of peaks at boundaries for this protein set. They all have between 5 and 15% overlap with boundaries, a similar genome-wide distribution to CTCF (Fig. S6e).

We investigated the presence of other proteins within the type I and type IV boundaries. We found that 86% and 71% of type I and type IV boundaries are occupied by other proteins in addition to CTCF, respectively (Fig. S6f). Type I and type IV boundaries, with CTCF, are also occupied by REST, ZNF316, SMC3, RAD21, ZNF143 and EMSY, as observed for boundaries genome-wide (Fig. 4b-c, S6g). 73% of type I boundaries without CTCF, are occupied by ZNF316, while in type IV boundaries this protein is found in 64% of the boundaries. Boundaries type I and IV without CTCF, are also occupied by EMSY and ATF2 (Fig. 4d-e, S6h).

Enrichment analysis of CTCF, ZNF316, EMSY and REST showed that these proteins are more enriched at type I boundaries (Fig. 4f). Type I and type IV boundaries without CTCF, are enriched in ZNF316 followed by EMSY and present little REST (Fig. 4g).

To determine if the boundaries might have different protein combinations, we measured the presence of CTCF, REST, ZNF316 and EMSY at the boundaries (Fig. 4h). The four proteins are present in 62% of type I boundaries. This proportion is lower at type IV boundaries (47%). Among the boundaries without CTCF, the most representative combinatory is ZNF316 and EMSY in type I and IV boundaries. 24 and 32% of type I and type IV boundaries respectively, do not have CTCF, REST, EMSY or ZNF316. Other proteins enriched at boundaries like ATF2, ATF4 and MAFF may occupy these boundaries (Fig. S6h).

Next, we analyzed the co-occupation between proteins in the same genomic regions within boundaries. As expected, CTCF co-localized with the cohesin complex subunits, and we found that CTCF and cohesin co-localize with REST very frequently. Interestingly, ZNF316 co-localizes primarily with EMSY without other factors (Fig. 4i). Given that CTCF’s contribution to boundary strength lies in the orientation of its DNA-binding motifs, we also explored the orientation for the REST and ZNF316. First, we determined the number of the ChIP-seq peaks at the boundaries (Fig. S7a). The number of peaks for these factors at boundaries is variable, ranging from 1 to 12 peaks per boundary, as we observed for CTCF. Then we identified the DNA-binding motifs for REST and ZNF316 (see methods) and their orientation. As for CTCF, the localization of the motifs was explored in windows of 10kb across the 50kb boundaries. The arrangement of motifs for ZNF316 and REST, showed no apparent preference (Fig. S7b). We then analyzed these proteins occupancy at both boundaries insulating a domain. We found that in many instances (more than 60%), ZNF316 and REST occupy both boundaries forming a type I-I domain. A lesser proportion was found for less insulated type IV-IV domain boundaries (Fig. S7c).

To explore whether REST, ZNF316 and EMSY are occupying loop anchors together with CTCF, we used previously reported loops in K562 cell line (9). As expected, CTCF is present in most loop anchor regions (84%). REST, SMC3, RAD21, and ZNF143 proteins are also present in more than 60% of these genomic regions. However, ZNF316 is only found at around 40% of loop anchors and EMSY in less than 40%. Thus, ZNF316 and EMSY are preferentially enriched at TAD boundaries, with ZNF316 occupying more than 80% of boundaries (Fig. S7d).

ZNF316 function has been described as a transcription factor (52,53). EMSY has been described as a transcriptional repressor interacting with BRCA2 that is recruited to chromatin as it does not have a DNA-binding domain (54–57). Our results suggest that ZNF316 (in some instances together with EMSY) could also structure domains in the presence or absence of CTCF.

### High accessibility boundaries are conserved across different cell types

To investigate if boundary accessibility relates to specific gene functions and boundary conservation, we first classified domains according to their boundaries’ accessibility. We considered domains in which both boundaries are type I, II, III or IV, as well as all possible combinations and identified their gene content. We focused on the proportion of 22% domains have type I boundaries at both sides and the 7% exhibit type IV boundaries (Fig. S8a).

Domains with start and end type I boundaries (I-I), would represent the most stable domains since these boundaries isolate contacts more efficiently. On the contrary, we suppose that domains with start and end type IV boundaries (IV-IV) could be more dynamic. To explore the variety of genes located inside domains I-I and domains IV-IV, we identified the genes inside these domains, measured their expression levels, and determined their function by gene ontology. The expressed genes in domains I-I are involved in essential and constitutive cellular functions like DNA packing complex, chromatin, regulation of transcription and signaling by G-protein-coupled receptors. Interestingly, also there are cell type specific functions, for example hematopoiesis and immune system (Fig. S8b). An example of this domain type houses one of the histone gene clusters and the *BTN* gene family, a subset of the immunoglobulins, all of which are highly expressed in K562s (Fig. S8f). In contrast, another example of an I-I domain that contains genes that are not expressed in K562 cells, harbors the olfactory receptor genes whose expression is tightly regulated during development (Fig. S8c-f). The genes inside domains IV-IV that are transcribed display general ontological categories, and the non-expressed genes are for example genes whose function are related to epidermis and tissue development (Fig. S8d-e). The developmental gene *FOXP2* is in a domain with type IV boundaries at each side and it is not expressed in K562 cells (Fig. S8g). We also measured the size of the different domain types and found that domains with type IV-IV boundaries are larger than type I-I domains (Fig. S8h).

Next, we assessed domain conservation across different human cell types. Several reports have shown that domain boundaries are conserved between cell types and species (1,54,55). However, these vary greatly depending on the resolution of the Hi-C matrices (60). To explore the conservation level of TADs with different accessibility degrees, we used published Hi-C data from different cell types and called domains and boundaries. We identified the conserved boundaries across different cell lines including GM12878, HCT116, HMEC, HUVEC, IMR90, NHEK and K562, and classified them according to their accessibility type in K562 cells (Fig. 5a). We found that 54% of conserved boundaries are highly accessible type I boundaries while only 19% are type IV (Fig. 5b).

**Figure 5.**
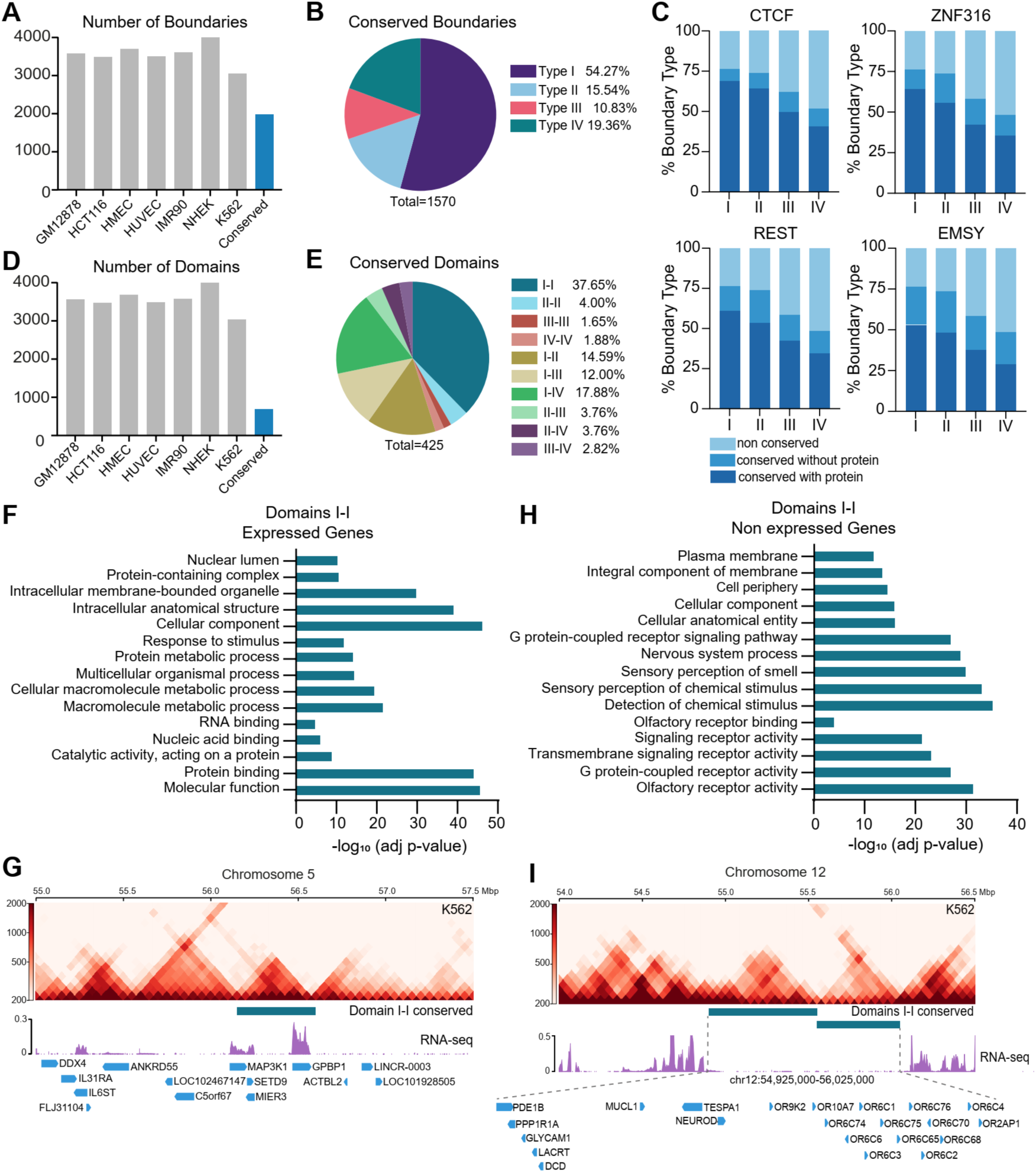
Highly accessible boundaries are conserved across human cell types. (A) Number of domain boundaries in different human cell types and boundaries identified as conserved in all data sets. The boundaries conserved between cell types correspond to 1985 boundaries. (B) From the conserved boundaries 1570 boundaries correspond to the ones classified based on their accessibility in our experiments. 54% of them are type I boundaries. (C) CTCF, ZNF316, REST, and EMSY occupancy at conserved boundaries. (D) Conserved domains across human cell types. (E) Classification of conserved domains according to their boundaries’ type. (F) GO analysis of the expressed genes within I-I conserved domains (G) An example of a conserved I-I domain containing the *MAP3KI* and *GPBP1* gene. The Hi-C matrix and RNA-seq tracks correspond to the K562 cell line. (H) GO analysis of non-expressed genes within conserved I-I domains. (I) An example of a conserved I-I domain harboring transcriptionally inactive genes in K562 cells. In this region, there is a cluster of genes that encode for olfactory receptors. The Hi-C matrix and RNA-seq tracks correspond to the K562 cells.

Then we inferred protein occupancy at conserved boundaries and found that ZNF316, EMSY and REST are occupying more than 50% of type I and II boundaries together with CTCF (Fig. 5c). These results suggest that these factors could be participating in boundary formation also in other cell types.

Next, we analyzed domain conservation considering both boundaries and found 684 conserved domains across the studied cell lines (Fig. 5d). 37.6% of conserved domains correspond to I-I domains with type I boundaries on either side and only 1.8% correspond to IV-IV domains with type IV boundaries on either side (Fig. 5e).

Then we explored the gene content of conserved I-I domains. Gene ontology enrichment analysis shows that these domains either contain highly expressed constitutive genes such the histone clusters and the kinase gene *MAP3K1* (Fig. S8f and 5f-g) or non-expressed genes that require tight gene expression regulation such as Olfactory Receptors coding genes (Fig. 5h-i).

These results suggest that domains with highly accessible and robust boundaries are the most conserved between different cell types and isolate highly active transcriptional units of constitutive genes or genes that need strict gene expression regulation in certain tissues and that these structures are preserved in different cells even if they are not being expressed.

### ZNF316 and EMSY participate in the formation and maintenance of domain boundaries

Our data suggest that ZNF316 might have a function in boundary formation as it is present in 76% of the boundaries detected (Fig. 6a). To explore if ZNF316 per se can participate in boundary formation, we selected a high accessibility boundary without CTCF and designed RNA guides to eliminate its DNA-binding site by CRISPR-Cas9 technology. We generated a homozygous mutant with a 612bp deletion containing the ZNF316 DNA-binding site. Our edition design also includes a ChIP-seq peak for EMSY that overlaps with ZNF316 occupancy (Fig. 6b, S9a,b). Next, we measured chromatin interactions using Hi-C. The Hi-C interaction matrix shows that the edited boundary does not disappear. However, inter- domain contacts that cross the edited boundary are gained. Also, the mutant exhibits a new interaction stripe absent in the control (Fig. 6c). The differential matrix shows that when the ZNF316 DNA-binding site is deleted at the selected boundary, there is a general increase in contact frequency (Fig. S9c). The TAD-separation score in the plotted region shows the affection in the edited boundary, as the control has a more robust score than the mutant (Fig. 6d-e). In contrast, the TAD-separation scores at a control region upstream of the edited area on the same chromosome behave similarly in both the control and mutant cells (Fig. S9d). We next quantified the contacts made from the edited region in the WT and mutant cells and found a reduction in contacts arising from the edited region in the mutant compared to the WT (Fig. S9e above) in contrast to two control regions (one upstream of the edited genomic region and one in a different chromosome), from which the number of contacts is equivalent between the WT and mutant cells (Fig. S9e below).

**Figure 6.**
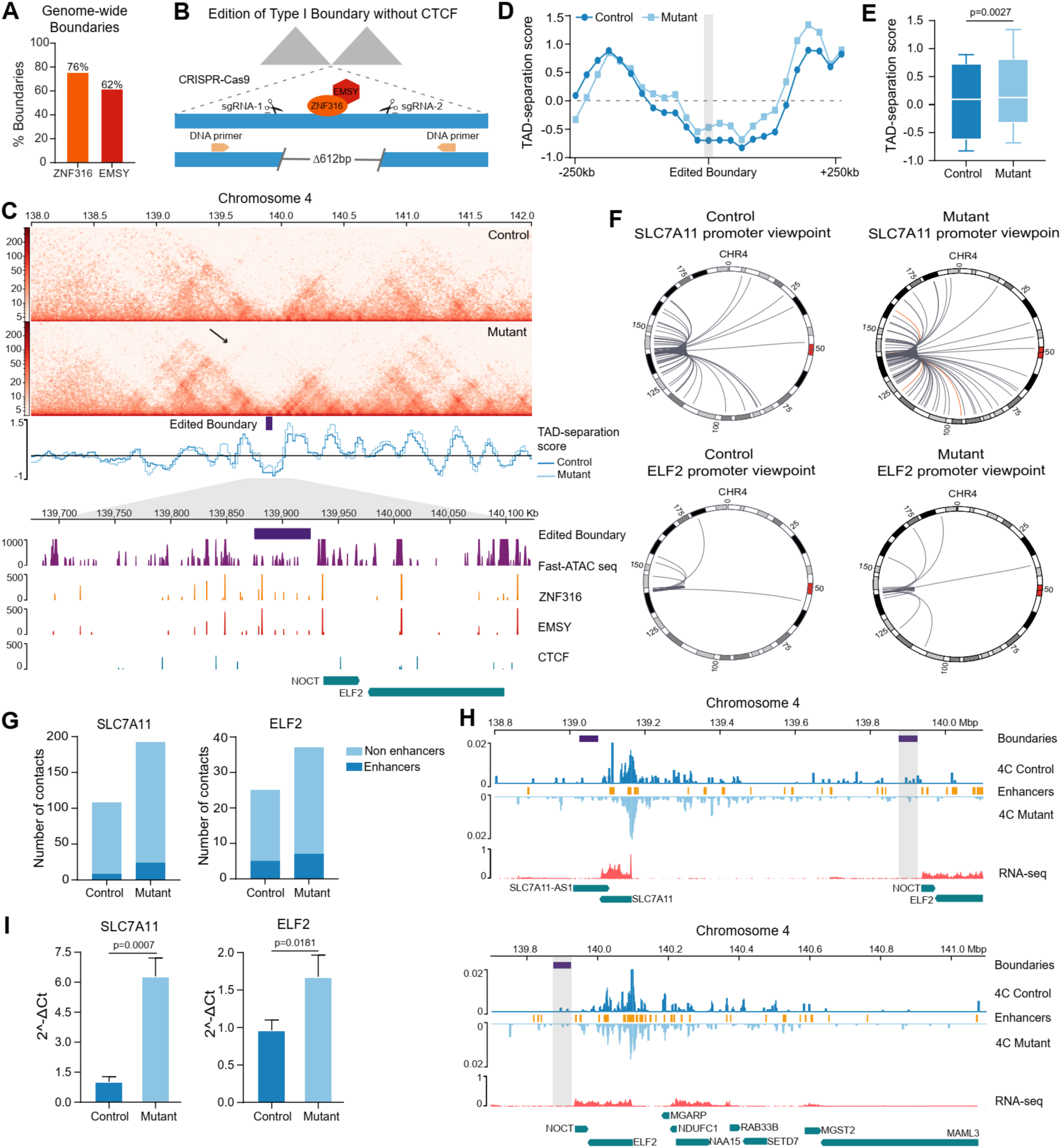
ZNF316 and EMSY block chromatin contacts across domain boundaries. (A) Proportion of boundaries with at least one ChIP-seq peak of ZNF316 and EMSY. (B) Diagram of the deletion performed of a region inside a type I boundary without CTCF, that contains a ChIP-seq peak of ZNF316 and EMSY. We applied CRISPR-Cas9 system and used two sgRNAs. The deletions were evaluated with a set of primers to amplify a region of 1kb in wild type (WT) K562 cell line (yellow arrows). (C) Hi-C matrices at 20kb resolution for the control and edited cells. The edited boundary is represented by a blue box (chr4:139870000-139890000). The arrow on the mutant matrix remarks a gain of inter-domain contacts. (D) TAD-separation score for each 20kb window from the Hi-C of the control and mutant cells. An area of +/- 270kb around the edited boundary is covered in the graph. (E) Box plot of the TAD-separation scores plotted in (D). The Wilcoxon test showed a significant statistical difference. (F) Intra-chromosomal interactions of two promoters surrounding the edited boundary. Above, circus plot showing the interactions from the *SLC7A11* gene promoter located upstream of the deletion site. Below, circus plot showing the interactions from the *ELF2* gene promoter located downstream of the deletion site. The orange arcs represent interactions with enhancers. Interactions in the WT and mutant cells are shown (G) Total number of intra-chromosomal interactions from the *SLC7A11* and *ELF2* gene promoters in WT and mutant cells. (H) Virtual 4C plots from the *SLC7A11* and *ELF2* gene promoters in WT and mutant cells. The modified boundary by CRISPR-Cas9 is marked in grey (chr4:139870000-139890000). (I) *SLC7A11* and *ELF2* gene expression measured by RT-qPCR in WT and mutant cells. The unpaired t-test found a statistically significant difference.

Next, we measured the contact number from gene promoters of genes flanking the edited boundary, and contacts are more numerous in the mutant than in the WT. For example, the *SLC7A11* (Solute Carrier Family 7 Member 11) and *ELF2* (E74 Like ETS Transcription Factor 2) genes located upstream and downstream of the altered boundary, respectively, present a gain in interactions in the mutant in contrast with the control (Fig. 6f-g). By performing a virtual 4C, the interaction profile shows increased contacts in the areas around the promoter, and some of these interactions are made with enhancers (Fig. 6h). We quantified the expression levels of these genes and observed in increased expression in the mutant compared to the control (Fig. 6i). Other genes surrounding the edited boundary also display more contacts in the mutant versus the control cells in contrast to a non-edited genomic region used as a control (Fig S9f). Our results show that ZNF316 (together with EMSY) has an insulating function at boundaries without CTCF blocking interactions across domains and could also block the extension of interaction stripes.

## DISCUSSION

Domains are essential for organizing the genome in chromatin neighborhoods and regulating gene expression in time and space. Domain boundaries are critical regions that delimit and constrain chromatin contacts. Many reports have demonstrated that genetic boundary alterations can have a functional impact on gene expression regulation by either forming or restricting chromatin contacts between regulatory elements such as enhancers and promoters (3,57).

In this work, we experimentally combined chromatin accessibility and Hi-C, to identify and classify domain boundaries according to their accessibility as a proxy for protein occupancy, to then describe in detail their genomic features. To do so, we modified the restriction enzyme incubation time points in the Hi-C assay, to gradually digest chromatin based on its accessibility. Finally, we identified TADs and classified the boundaries into four accessibility types. To give a detailed description of the genomic features of domain boundaries, we focused on the most accessible boundaries (Type I) and contrasted them with the least accessible boundaries (Type IV). Type I boundaries are occupied by more proteins and result in better insulators of chromatin contacts compared to type IV boundaries, with lower TAD-separation scores and fewer inter-TAD contacts crossing the boundary.

Although CTCF function in domain boundary formation and loop extrusion has been extensively documented (19,21,22,25,58–63), some reports show that CTCF depletion does not completely affect domain formation and provides evidence of domain boundaries that are CTCF-independent (11,26,27,64). Even though most of our detected boundaries have CTCF, we found that 11 and 26% of type I and type IV boundaries respectively, are not occupied by CTCF. Interestingly, the TAD-separation score and the number of interactions blocked by type I boundaries with or without CTCF are equivalent, suggesting other factors contribute to boundary insulation properties besides CTCF.

The convergent CTCF motif orientation at loop or domain anchors is necessary to efficiently block the cohesin complex according to the loop extrusion model (21,23,24,47,48,65). Boundaries with the highest accessibility showed this divergent pattern of CTCF binding motifs, in contrast with the lowest accessibility boundaries, suggesting that this arrangement is relevant to the insulating properties of the boundary. Interestingly, we also found that the conservation score of the CTCF DNA-binding site in type I boundaries is higher than the motif found in type IV boundaries. This might reflect CTCF binding affinity or residence time at the more conserved motif giving the boundary more stability.

To characterize which other proteins occupy boundaries we explored ChIP-seq data in K562 cells and found that REST, ZNF316 and EMSY occupy >60% of the boundaries with CTCF. Our data also found SMC3 and RAD21 and other previously reported proteins occupying boundary elements (18,66). CTCF and Cohesin commonly co-localize with REST, and ZNF316 often co-localizes with EMSY at boundaries, constituting two independent protein modules. Interestingly, ZNF316 is present in 70% of boundaries without CTCF.

Finally, we functionally tested ZNF316 and EMSY function at boundaries by CRISPR-Cas9 mediated edition of ZNF316 DNA-binding site at a selected highly accessible boundary without CTCF. EMSY does not bind DNA directly, so ZNF316 might recruit it at domain boundaries. The edition significantly increases intra-domain contacts in contrast to the non-edited control. Also, there is an increase in the inter-domain contacts from the promoters surrounding the edited boundary, some with enhancers. For some of these genes, we detected an upregulation in gene expression in the mutant compared to the control. Our results show that ZNF316 (together with EMSY) contribute to domain boundary formation without CTCF.

## DATA AVAILABILITY

Data produced in this study will be available under GSE268028 and GSE268108 accession numbers, once the manuscript is accepted. The GEO accession numbers for the published ChIP-seq data sets (53), RNA-seq data sets (53), ATAC-seq data sets (34) and Hi-C data sets (9) used are listed in Supplemental Table S4.

## ACKNOWLEDGMENTS

This paper is part of the requirements to obtain a doctoral degree under the Postgrado en Ciencias Biológicas, UNAM for K.J.-L. We thank the Molecular Biology Unit at IFC and Dr. Rosa Rebollar from la RAI for her help with sample sonication. We thank Juan Carlos Gómora for reading and commenting of the manuscript. We thank Dr. César Poot-Hernández and Ms. Carlos-Peralta for their help with computer servers’ management, and software installation and data upload to GEO. We thank Dr. Felix Recillas-Targa and Ms. Georgina Guerrero for their assistance during the first stages of this projectߣs development. We thank the Supercomputing Unit at LANCIS, in particular R. García-Herrera for access to the computer cluster.

