## Supplemental_figures for "Chromatin accessibility classification of TAD boundaries discloses new architectural proteins"

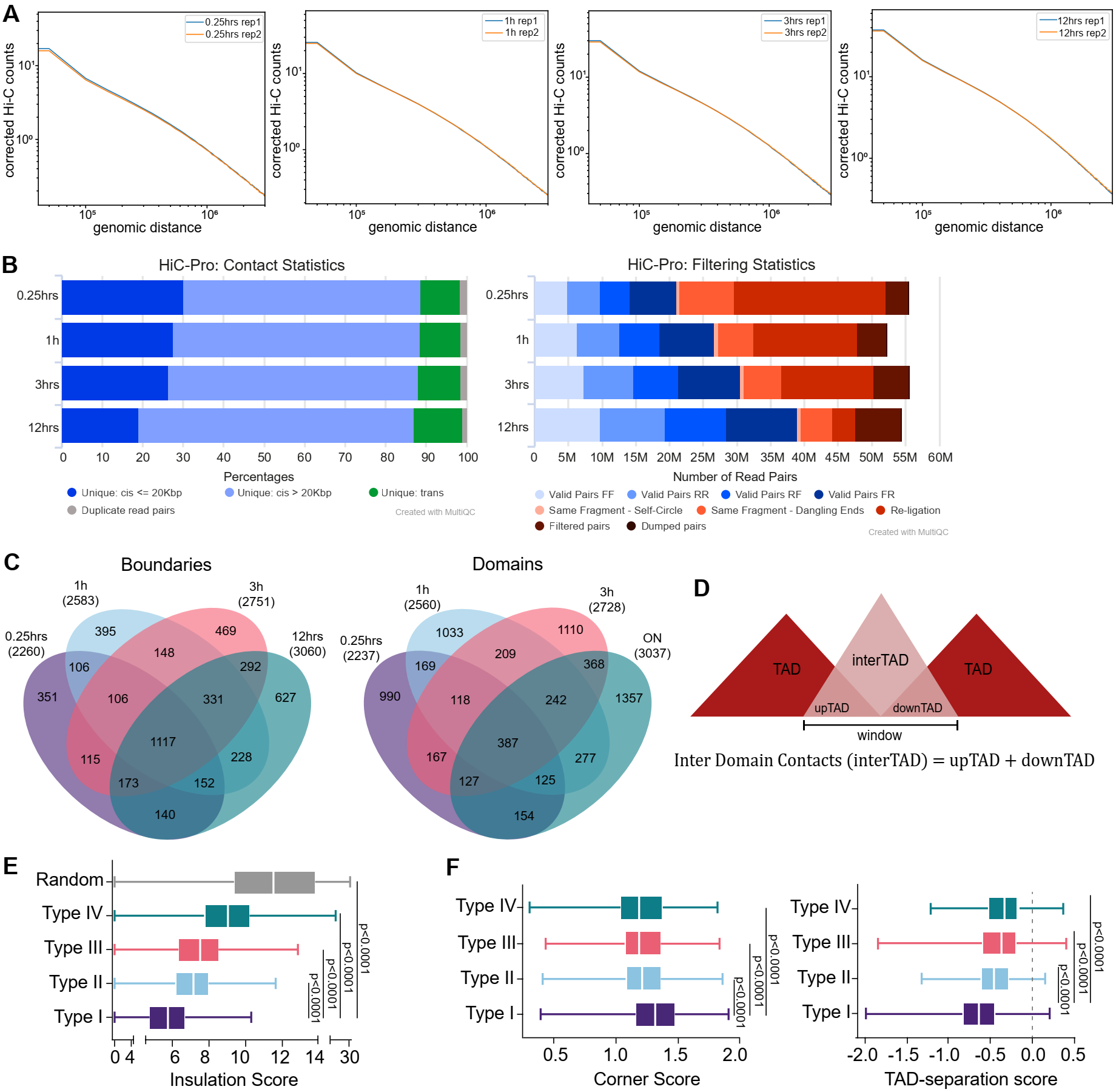

**Supplemental Figure S1. Hi-C libraries quality controls and boundary classification.** (A) Hi-C corrected counts enrichment at different genomic distances plotted for all replicates. The results reflect the proportion of long-range and short-range contacts. The Hi-C counts at different distance ranges displays the same distribution between replicates indicating equivalent *cis* decay profiles. (B) The left graph shows the number of valid pairs obtained (R: reverse fragments and F: forward fragments) and all the categories of invalid pairs. On the right the graph shows the percentage of valid pairs in both *cis* and *trans*. All experiments were adjusted to the number of valid pairs obtained in 0.25 hours. (C) In order to describe the genomic features within the boundaries, we choose the following groups of boundaries: 1117 detected in all Hi-C assays (Type I boundaries), 331 detected at 1,3 and 12 hours (Type II boundaries), 292 detected at 3 and 12 hours (Type III boundaries) and 627 detected at 12 hours (Type IV boundaries). (D) The Inter-TAD was determined by calculating as described (see methods). (E) Insulation scores of the boundary types. Kruskal-Wallis test was applied followed by Dunn´s post hoc test. (F) For boundaries mapped at 5kb resolution, Corner Scores and TAD-separation scores are shown for all boundary types. Kruskal-Wallis test was applied followed by Dunn´s post hoc test.

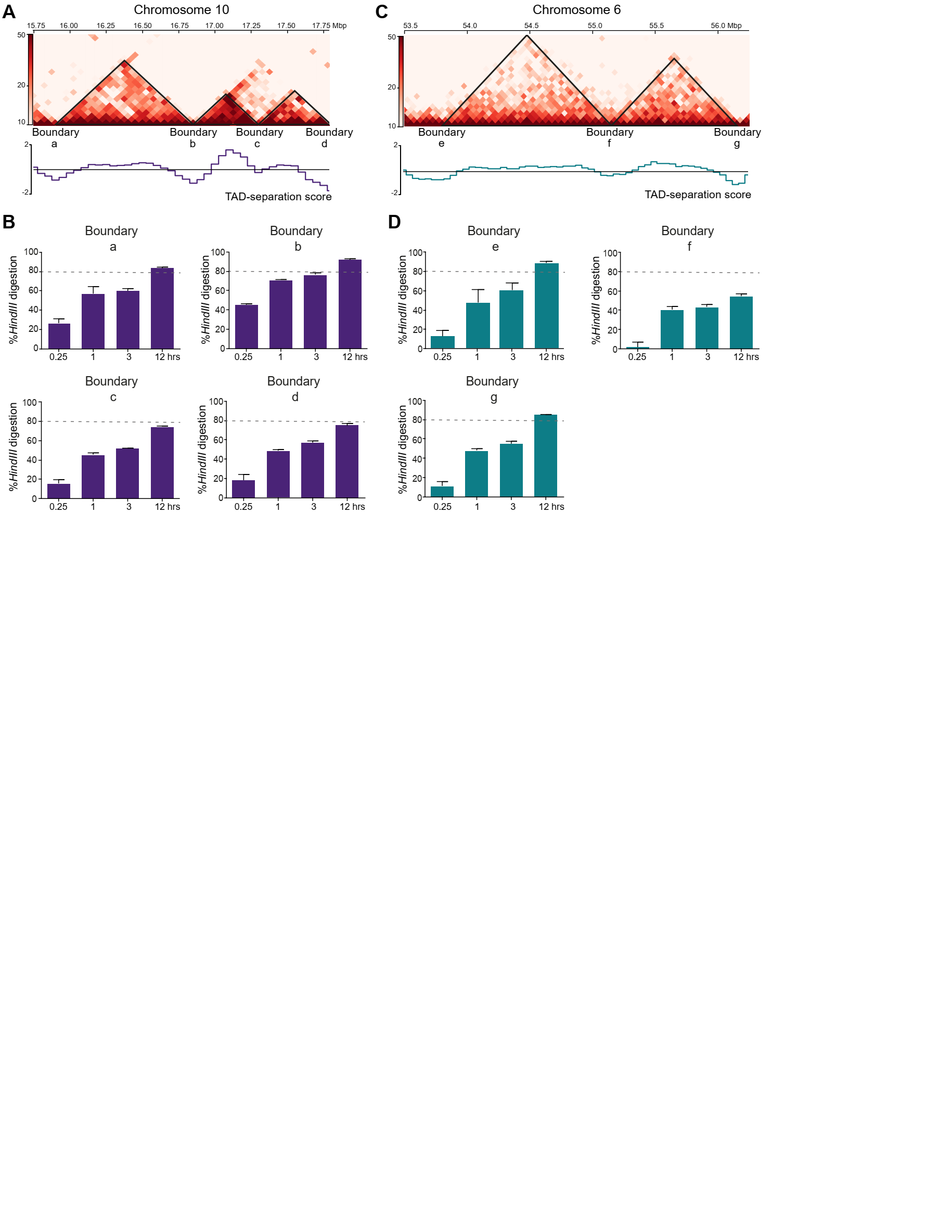

**Supplemental Figure S2. Type I boundaries are more accessible than type IV measured by qPCR.** (A) Hi-C matrix with examples of highly accessible type I boundaries on chromosome 10. (B) Digestion percentages of the amplicons containing type I boundaries indicated in A in chromosome 10. The boundary with higher restriction is boundary b, it showed more than 40% of digestion since 0.25 hours, n=3. (C) Hi-C matrix with low accessibility type IV boundaries on chromosome 6. (D) Digestion percentages of the amplicons containing the boundaries indicated in C in chromosome 6. For the three evaluated boundaries, the percent of restriction is below of 20% at 0.25 hours, n=3.

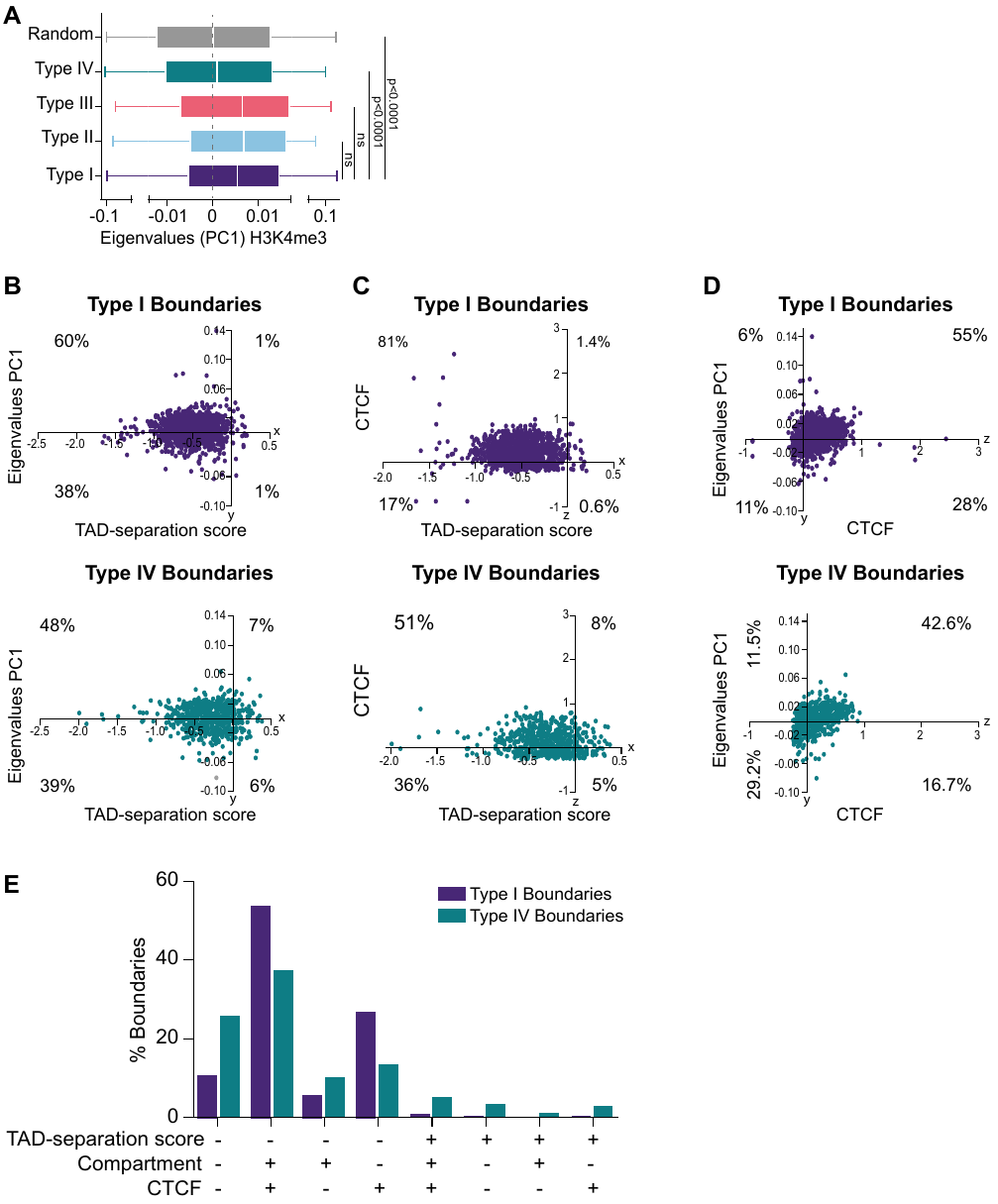

**Supplemental Figure S3. Correlation between TAD-separation scores, Compartment assignment (A/B) and CTCF occupancy in the different type of boundaries.** (A) PC1 eigenvalues from PCA analysis to determine the (A/B) compartment using H3K4me3 active histone mark as a seed to assign the A compartment (positive values). Kruskal-Wallis test followed by Dunn´s post hoc test was applied. (B, C, D) Graphs with three axes, x: TAD-separation score, y: PC1 eigenvalues (H3K27ac) and z: CTCF average signal. The panels with blue dots correspond to type I boundaries, the graphs with green dots represent the type IV boundaries. (B) 2D view graph for TAD-separation score and PC1 values. (C) 2D view graph for TAD-separation score and CTCF. (D) 2D view graph for PC1 values and CTCF. (E) Percent of type I and type IV boundaries, grouped according to the values (positive or negative) for TAD-separation score, PC1 eigenvalue (H3K27me3) and CTCF average signal.

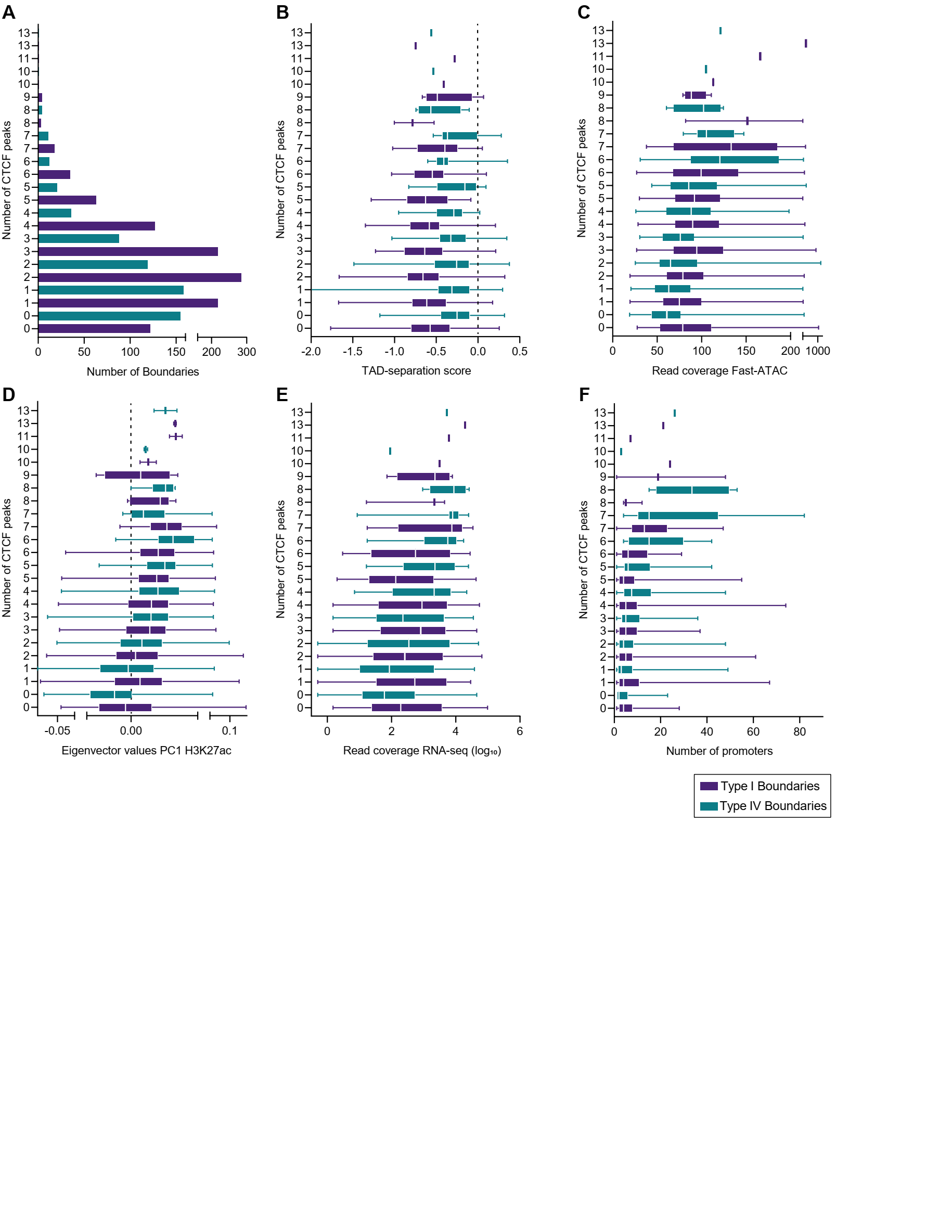

**Supplemental Figure S4. Boundary features based on the number of CTCF peaks occupying the boundary.** (A) Number of CTCF ChIP-seq peaks in type I and type IV boundaries. (B) TAD-separation scores for type I and type IV boundaries with different number of CTCF peaks. (C) Fast-ATAC seq read coverage at type I and type IV boundaries with different number of CTCF peaks. (D) PC1 eigenvalues for type I and type IV boundaries with different number of CTCF peaks using H3K27ac histone mark as the seed for compartment A assignment (positive values). (E) RNA-seq read coverage at type I and type IV boundaries with different number of CTCF peaks. (F) Number of promoters at type I and type IV boundaries with different number of CTCF peaks.

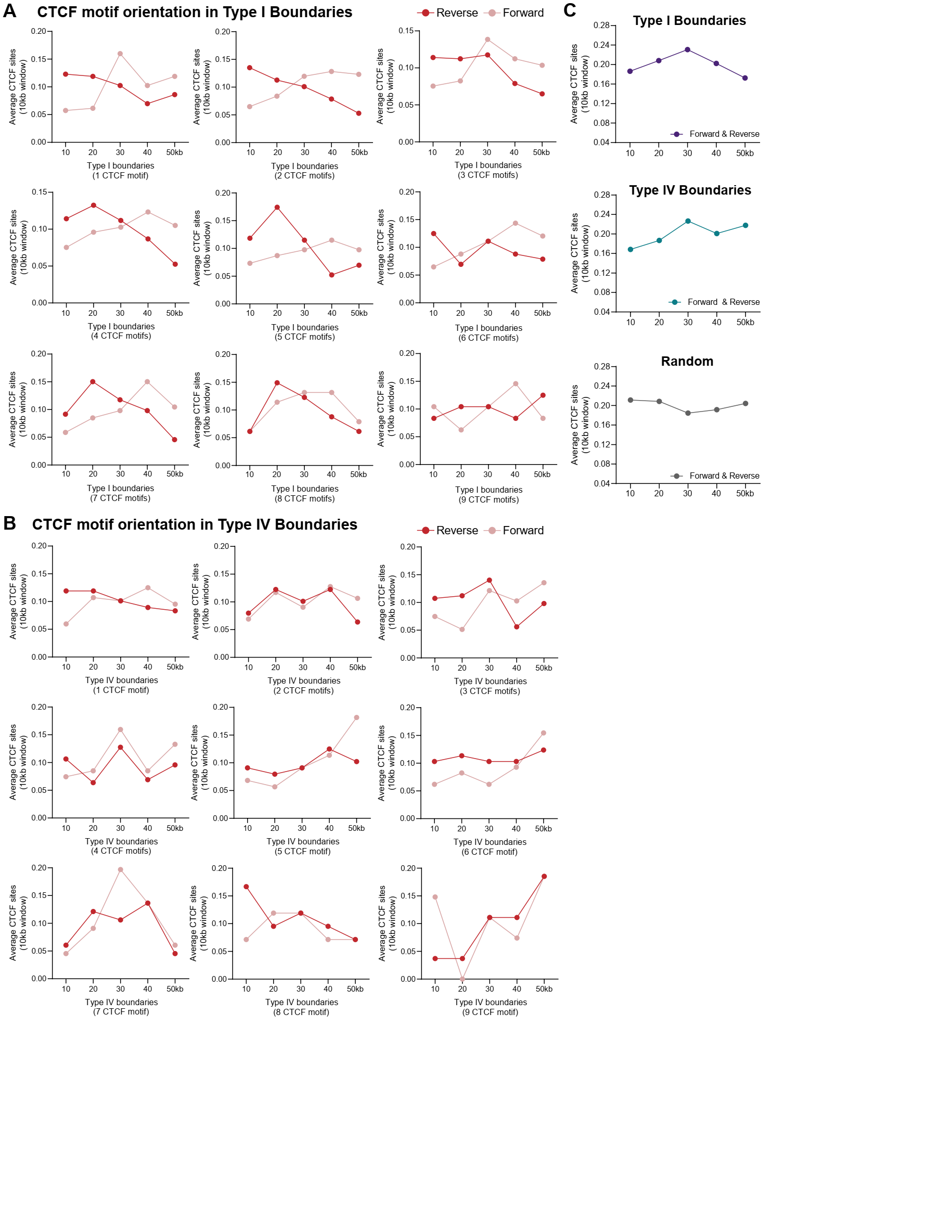

**Supplemental Figure S5. CTCF DNA binding motifs orientation at type I and IV boundaries with different number of CTCF motifs.** (A, B) CTCF DNA binding motif orientation analysis. Each graph corresponds to the boundaries grouped by the number of CTCF motifs inside the boundaries. Plotted is the average CTCF motif in forward and reverse orientation along the 50kb boundaries, each 10kb window. The red dots represent the reverse orientation motif, while the pink dots correspond to the forward orientation motif. (A) Type I boundaries. (B) Type IV boundaries (C) Average orientation of all CTCF DNA binding motifs (reverse and forward) along the 50kb boundaries type I, type IV and random regions.

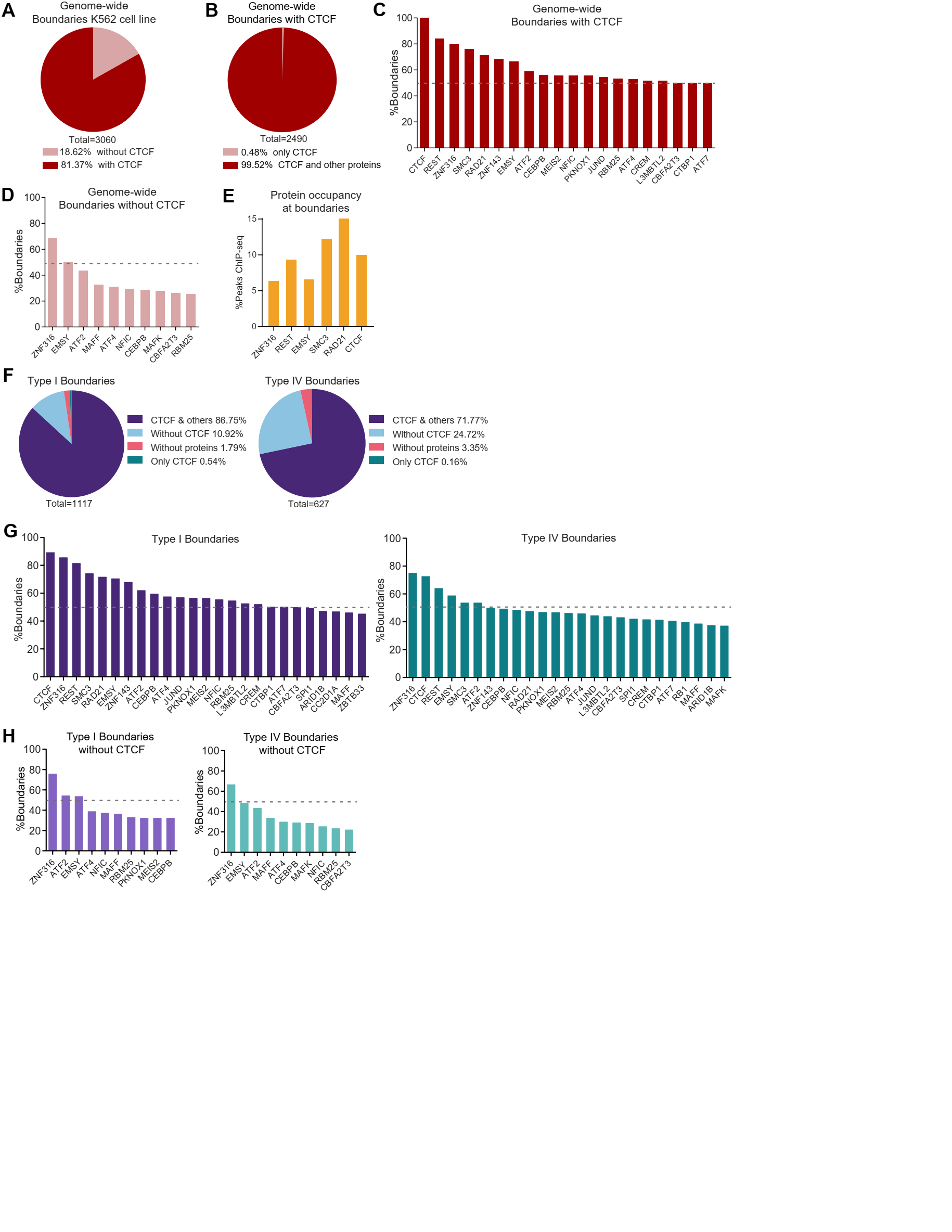

**Supplemental Figure S6. Boundary protein occupancy analysis for the 296 ChIP-seq data sets in K562 cell line.** (A) Percent of boundaries in K562 cell line with and without a CTCF ChIP-seq peak. (B) For the boundaries occupied by CTCF, fraction of boundaries that present only CTCF and fraction of boundaries with CTCF an also another protein from the 296 ChIP-seq database. (C) For the boundaries occupied by CTCF, each bar corresponds to the percentage of boundaries with other protein indicated in the x axis. The dotted line marks 50% of boundaries. Proteins occupying less than 50% of the boundaries are not shown (D) For the boundaries not occupied by CTCF. Each bar corresponds to the fraction of boundaries occupied by other protein from the 296 ChIP-seq database. The dotted line marks 50% of the boundaries. Proteins occupying less than 25% of boundaries are not shown. (E) For each protein on the x axis, percentage of peaks located at domain boundaries. As for CTCF, most proteins occupy many regions in the genome besides domain boundaries. (F) Boundaries type I and type IV classified by the presence of CTCF only, CTCF and other protein from the 296 protein database, absence of CTCF and boundaries without any protein. (G) For all type I and IV boundaries, each bar corresponds to the fraction of boundaries occupied with the specified protein on the x axis. The dotted line marks 50% of the boundaries. The first 25 proteins are shown (H) For type I and IV boundaries without CTCF, each bar corresponds to the fraction of boundaries occupied by the protein indicated. The dotted line marks 50% of the boundaries. The first 10 proteins are shown.

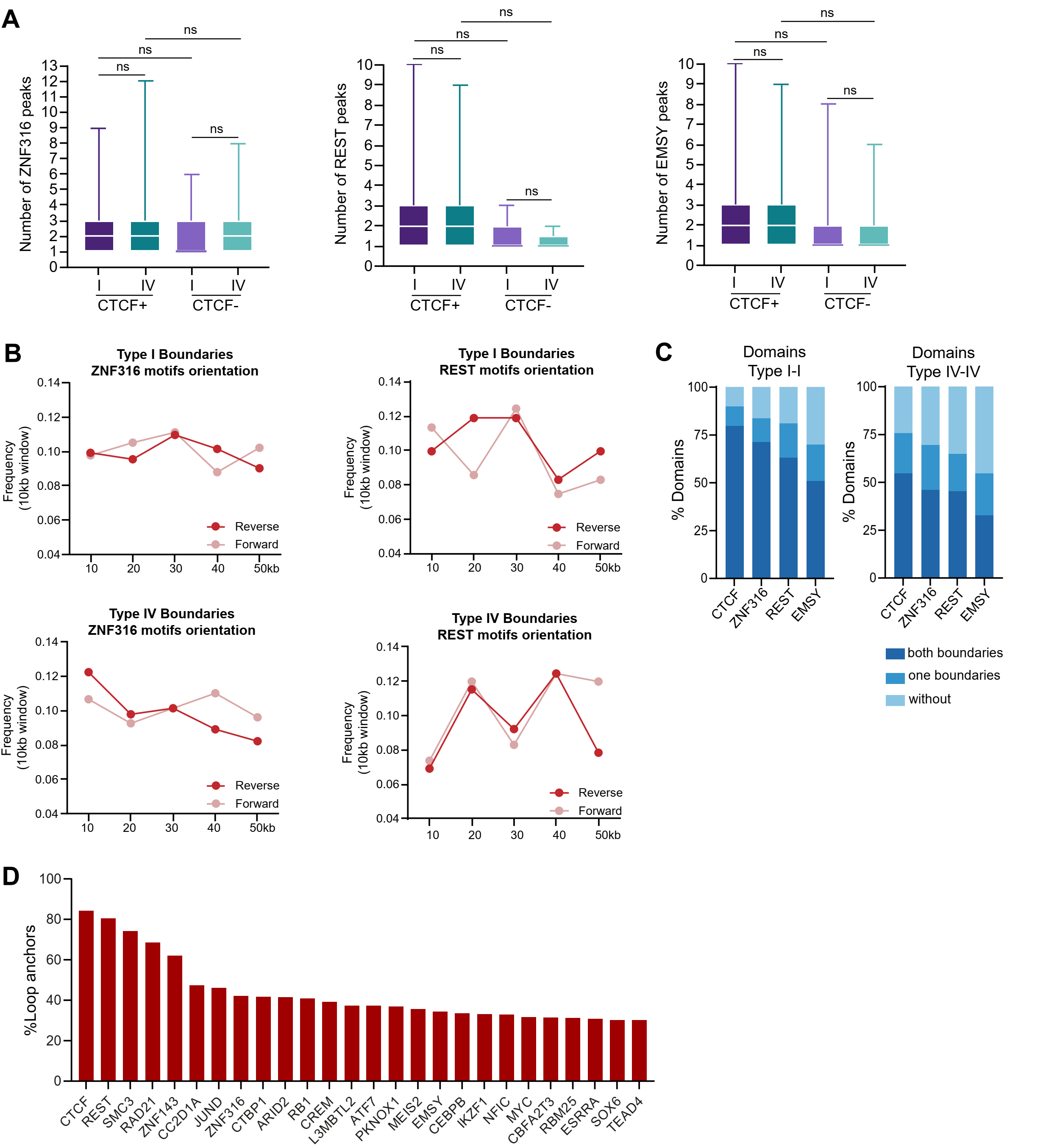

**Supplemental Figure S7. REST, ZNF316 and EMSY occupancy and DNA binding motifs orientation at domain boundaries.** (A) ChIP-seq peak number for REST, ZNF316 and EMSY at type I and type IV boundaries with (CTCF+) and without (CTCF-) CTCF. Kruskal-Wallis test followed by Dunn´s post hoc test was applied (ns, no significate). (B) ZNF316 and REST average motifs in forward and reverse orientation along the 50kb boundaries each 10kb windows. The red points represent the reverse orientation motif, while the pink points correspond to the forward orientation motif. (C) CTCF, REST, ZNF316 and EMSY occupancy in one or both boundaries of I-I and IV-IV domains. (D) Percentage of loop anchors occupied by proteins from the 296 ChIP-seq data sets in K562 cell line. Proteins occupying less than 30% of anchors are not shown.

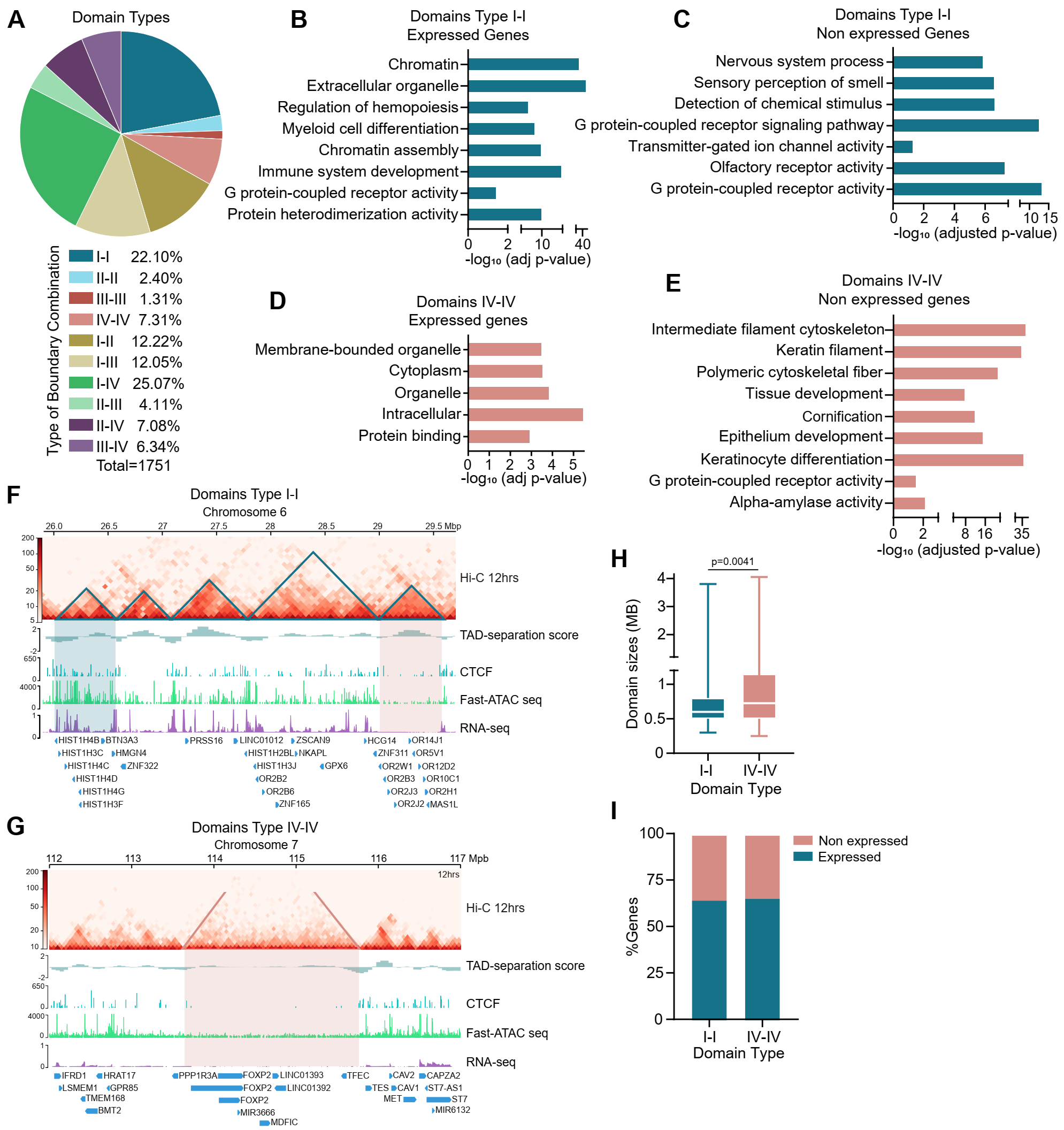

**Supplemental Figure S8.** **Genes inside domains with high accessibility are related to constitutive and cell type-specific functions.** (A) The total domains were classified based on their accessibility in the start and end boundaries. (B) The domains with both boundaries classified as type I (domains I-I), contained transcriptionally active genes. These genes are related to constitutive cell functions. (C) Transcriptional inactive genes were also present in the domains I-I. The gene ontology enrichment showed some categories unrelated to the K562 cell line, such as olfactory receptor genes. (D) The domains with both boundaries classified as type IV (domains IV-IV) that contained transcriptionally active genes present genes with general cellular functions. (E)The transcriptionally inactive genes inside IV-IV domains showed ontological categories related to epidermal tissues. (F) The Hi-C matrix shows a region in chromosome 6 with I-I domains. A group of transcribed histone genes is present in the first domain. On the other hand, the last domain contains olfactory receptor genes that are transcriptionally inactive. (G) The Hi-C matrix shows a region in chromosome 7 with IV-IV domains. An example of the developmental gene *FOXP2* is displayed. (H) Box and whiskers plot of domain sizes. Mann-Whitney statistical test was applied. (I) Percentage of expressed and non-expressed genes inside I-I and IV-IV domains.

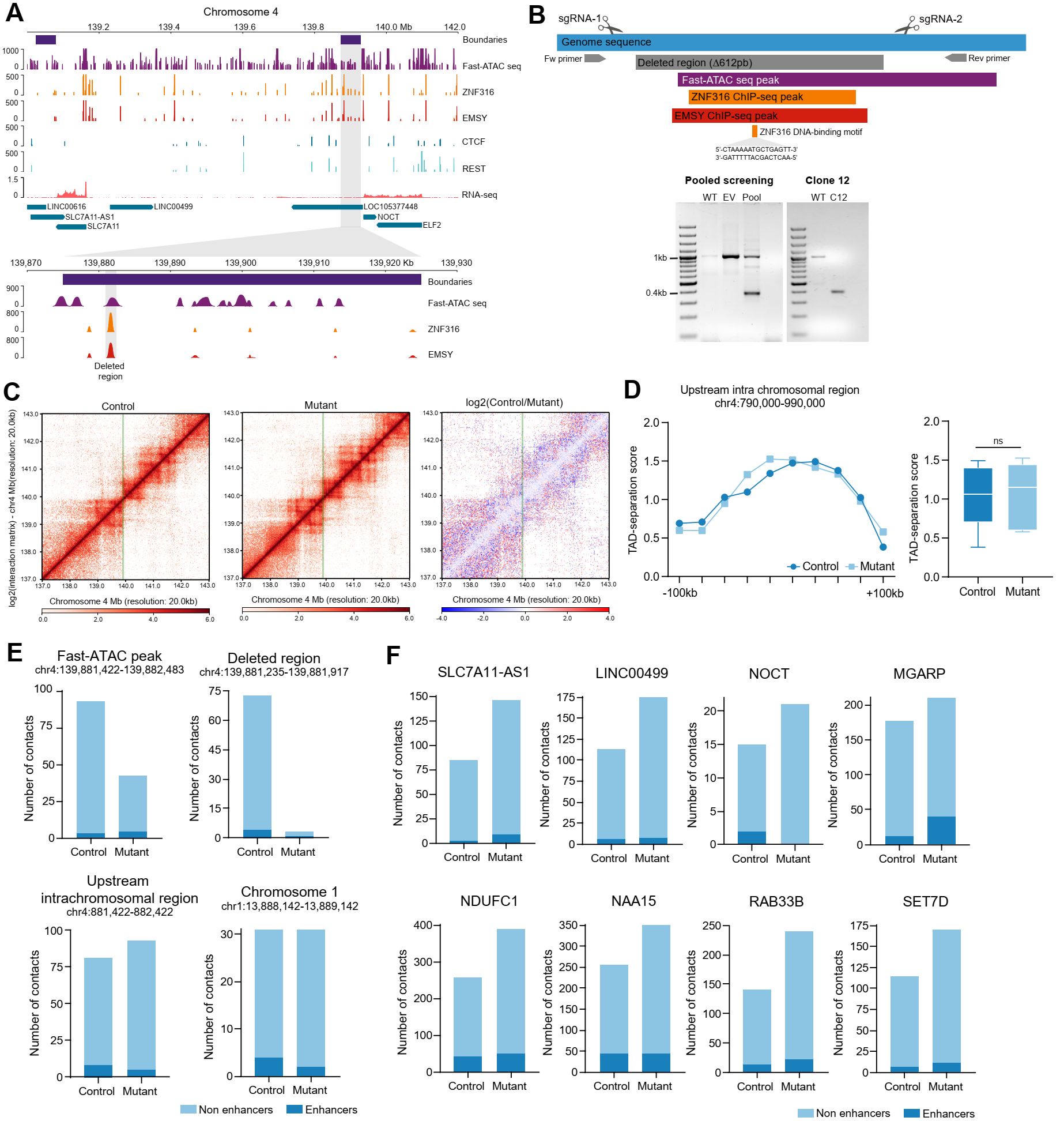

**Supplemental Figure S9. ZNF316 DNA binding motif deletion causes an increase in chromatin contacts from the surrounding gene promoters.** (A) The type I boundary selected to perform genetic edition of the ZNF316 DNA binding site. Hi-C matrix at 20kb resolution is shown. In the zoomed region the boundary is indicated in gray. ChIP seq signal for ZNF316, EMSY, CTCF and REST are presented. CTCF and REST are not occupying this boundary. The genes surrounding the boundary are indicated. (B) Boundary type I without CTCF edited by CRISPR-Cas9. Inside the boundary, we selected the area with higher accessibility and occupied by CTCF and EMSY (Peak of Fast-ATAC seq data). The sequence of the ZNF316 motif was confirmed. We designed two sgRNA to get a homogenous deletion. (C) Differential Hi-C matrix at 20kb resolution. The green line indicates the edited boundary. (D) TAD-separation scores for a control region upstream of the edited boundary. The scores were extracted for 20kb bins. A Wilcoxon test was applied, showing no statistically significant differences between the control and mutant. (E) Number of contacts from four different viewpoints in the control and mutant cells. The first viewpoint was the Fast-ATAC seq peak at the edited boundary. The second viewpoint corresponds to the exact deleted region inside the accessibility peak. As controls, a third viewpoint was a region located 138Mb upstream of the edited region. Finally, a fourth control region on a different chromosome was used to compare the number of contacts between the control and mutant cells. (F) Number of contacts from gene promoters surrounding the deleted region. The gene name is indicated above the plot.
